# High-throughput profiling of drug interactions in Gram-positive bacteria

**DOI:** 10.1101/2022.12.23.521747

**Authors:** Elisabetta Cacace, Vladislav Kim, Michael Knopp, Manuela Tietgen, Amber Brauer-Nikonow, Kemal Inecik, André Mateus, Alessio Milanese, Marita Torrissen Mårli, Karin Mitosch, Joel Selkrig, Ana Rita Brochado, Oscar P. Kuipers, Morten Kjos, Georg Zeller, Mikhail M. Savitski, Stephan Göttig, Wolfgang Huber, Athanasios Typas

## Abstract

Drug combinations present a powerful strategy to tackle antimicrobial resistance, but have not been systematically tested in many bacterial species. Here, we used an automated high-throughput setup to profile ∼ 8000 combinations between 65 antibacterial drugs in three Gram-positive species: the model species, *Bacillus subtilis* and two prominent pathogens, *Staphylococcus aureus* and *Streptococcus pneumoniae*. Thereby, we recapitulate previously known drug interactions, but also identify ten times more interactions than previously reported in the pathogen *S. aureus*, including two synergies that were also effective in multi-drug resistant clinical *S. aureus* isolates *in vitro* and *in vivo*. Interactions were largely species-specific and mostly synergistic for drugs targeting the same cellular process, as observed also for Gram-negative species^1^. Yet, the dominating synergies are clearly distinct between Gram-negative and Gram-positive species, and are driven by different bottlenecks in drug uptake and vulnerabilities of their cell surface structures. To further explore interactions of commonly prescribed non-antibiotic drugs with antibiotics, we tested 2728 of such combinations in *S. aureus*, detecting a plethora of unexpected antagonisms that could compromise the efficacy of antimicrobial treatments in the age of polypharmacy. We uncovered even more synergies than antagonisms, some of which we could demonstrate as effective combinations in vivo against multi-drug resistant clinical isolates. Among them, we showed that the antiaggregant ticagrelor interferes with purine metabolism and changes the surface charge of *S. aureus,* leading to strong synergies with cationic antibiotics. Overall, this exemplifies the untapped potential of approved non-antibacterial drugs to be repurposed as antibiotic adjuvants. All data can be browsed through an interactive interface (https://apps.embl.de/combact/).

## Introduction

Antibacterial agents have been used in combination since the dawn of the antibiotic era for different purposes: to achieve synergy (e.g. sulfamethoxazole-trimethoprim), to limit resistance (e.g. combinations of beta-lactams and beta-lactamase inhibitors, or antitubercular regimens), and/or to broaden the spectrum of action of anti-infective treatments (e.g. empiric treatments of sepsis)^2^. With antimicrobial resistance (AMR) posing a global threat to public health, which permeates all domains of modern medicine^3, 4^, the use of drug combinations to re-sensitize resistant strains has emerged as one of the promising means to bypass the stagnant drug discovery pipeline^5^.

Although a few antibacterial combinations are used in clinics, and screens for approved compounds as adjuvants for antibiotics have been increasingly conducted in the last decade^6–, 11^, the full potential of drug combinations for treating bacterial pathogens remains underexplored. This is because the combinatorial space is vast, and drug interactions are rare, and concentration-, drug-, time-, species- and even strain-specific^1, 12, 13^, making systematic testing necessary, yet highly demanding. As a result, drug interactions have not yet been systematically profiled in many clinically relevant bacterial species.

With the increase of polypharmacy^14^, antibiotics are often prescribed in combination with other drugs^15^. While pharmacokinetic interactions between antibiotics and non-antibiotic drugs are well-known for the host (e.g. dependencies on drug metabolism and excretion by the liver and the kidney)^16^, they are poorly characterized at the level of bacterial physiology.

We recently used automated platforms to systematically profile antibiotic interactions on three Gram-negative pathogens^1^. Here, we expanded this experimental and computational framework to generate a comprehensive resource of drug interactions in three Gram-positive bacterial species: the pathogens *Staphylococcus aureus* and *Streptococcus pneumoniae*, two of the most prominent antibiotic-resistant bacteria^3, 17^, and the model organism *Bacillus subtilis*. Compared to previous interaction studies in Gram-positive species^8, 10, 18^, this vastly increased the number of drugs, concentrations and strains tested. By probing all main classes of antibiotics, we could relate interaction outcomes to bacterial structural features, cellular network architecture, as well as to drug target conservation. Moreover, we profiled the interactions of antibiotics with a large panel of non-antibiotic drugs in *S. aureus* to investigate the impact of commonly administered medications on antibiotic efficacy. Thereby, we uncovered both strong synergies that remained effective against multidrug-resistant clinical *S. aureus* isolates, and widespread antagonisms that could compromise the efficacy of antibiotic treatments.

## Results

### An automated pipeline for high-throughput testing of drug combinations

We profiled 1891-2070 drug combinations in a 4 x 4 dose matrix (two-fold dilution gradient) in *Staphylococcus aureus*, *Streptococcus pneumoniae* and *Bacillus subtilis*. For *S. aureus*, two strains (*S. aureus* Newman and DSM 20231) were probed to assess within-species conservation (**Fig. 1a**, **Supplementary Table 1**). The drug panel (n = 65) included antibiotics (n = 57) used against infections with Gram-positive bacteria and belonging to all main classes and targeting different bacterial processes, and eight other bioactive molecules, such as antifungals, drugs with human targets and food additives – depicted as non-antibiotics (**Fig. 1a, Supplementary Table 1**).

**Figure 1.**
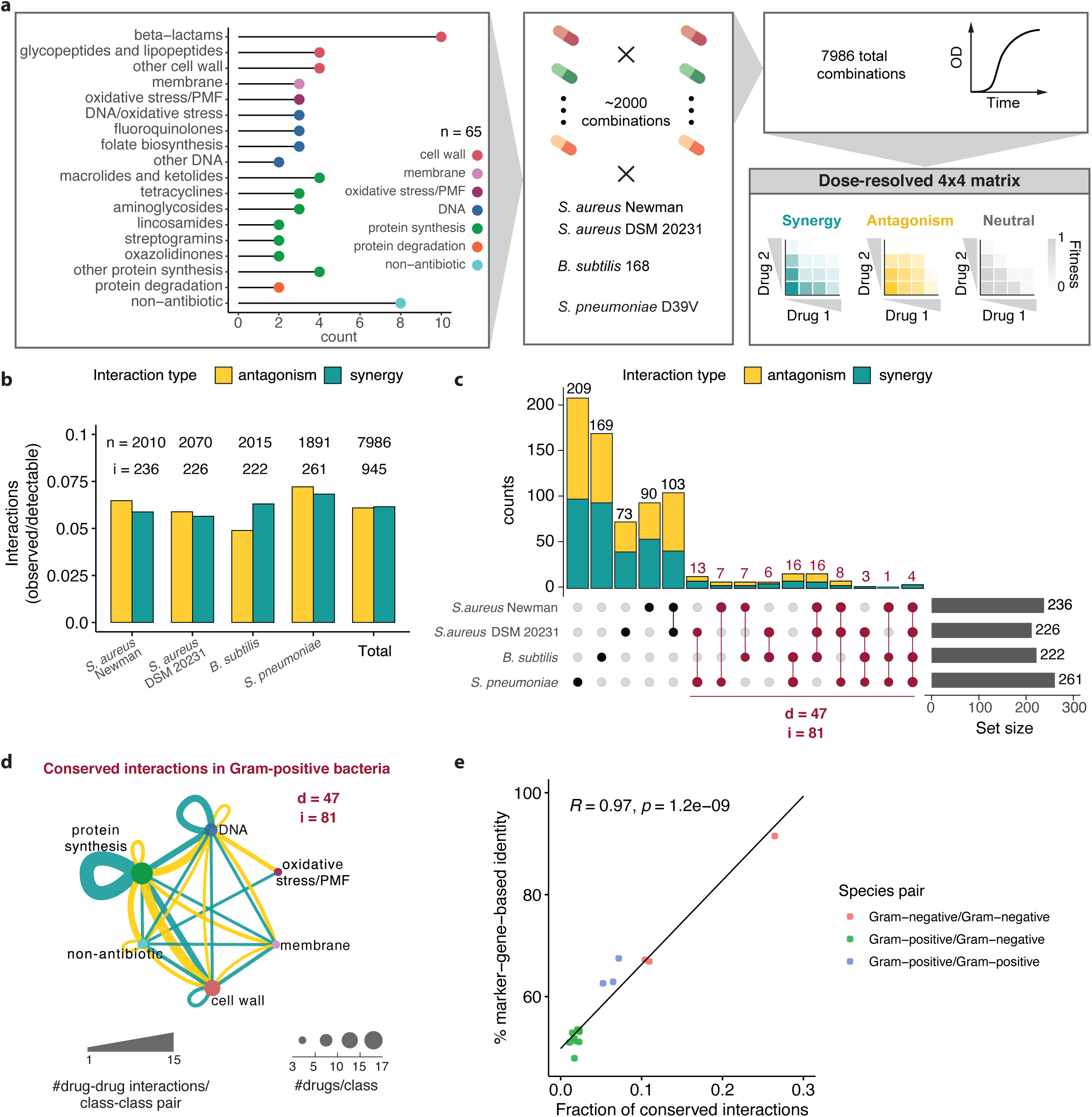
Drug-drug interactions are rare and species-specific in Gram-positive bacteria. **a**, Schematic representation of the high-throughput screen. Pairwise combinations of 65 drugs belonging to several chemical classes and targeting different cellular processes (**Supplementary Table 1)** were tested at three concentrations in three bacterial species: *S. aureus* (two strains), *B. subtilis* and *S. pneumoniae*. For each strain, 1891 to 2070 combinations were tested in broth microdilution in 384-well-plates, measuring OD_595nm_ over time. Normalised fitness values were calculated and used to obtain 4 x 4 checkerboards and assign interactions as synergistic, antagonistic or neutral (Methods**, Extended Data** Fig. 1**, Supplementary Table 2**). PMF = proton-motive force. **b**, Interaction abundance in each strain separately, and altogether. Synergy and antagonism frequencies are obtained dividing their absolute counts by the number of combinations for which the probed fitness space allows to detect synergy (fitness upon combination ≥ 0.1) or antagonism (fitness upon combination ≤ 0.9) discovery (Methods). Total numbers of combinations tested (n) and detected interactions (i) are shown for each set. **c**, Conservation of interactions among the four strains tested. All unique interactions detected in the screen (n = 725) are considered to calculate intersection sets between strains. The total number of interactions in each strain is indicated as set size, adding up to 945 total interactions in all strains. The 81 interactions (i), involving 47 drugs (d), are conserved across species (dark red). **d**, Network of conserved interactions between Gram-positive species. Drugs are grouped according to their targeted cellular process (**Supplementary Table 1**). Edge thickness is proportional to the number of drug-drug interactions for each class-class pair. Node size is proportional to the number of drugs in each class. Only drugs involved in this interaction set are considered (d = 47). Nodes are coloured according to the targeted cellular processes as in Fig. 1a. **e**, Drug interaction conservation between species recapitulates phylogeny. Pearson correlation between sequence identity (based on 40 conserved marker genes) and drug interaction conservation rate between pairs of species tested here and in a previous study^1^.

We measured growth in a broth microdilution format in microtiter plates using optical density (OD_595nm_) as a readout. Media and shaking conditions were different for each species (Methods). Drug concentrations were tailored after measuring minimal inhibitory concentrations (MICs) for each drug in the four strains, with the highest concentration corresponding to the MIC in most cases, and the intermediate and lowest concentration corresponding to half and a quarter of the highest concentration, respectively (Methods, **Supplementary Table 1**). We derived fitness values in the presence of single drugs and drug combinations, dividing single-time point OD_595nm_ values upon drug treatment by the corresponding values of no-drug controls at the same time point. This time point was selected according to the growth characteristics of each strain and aimed to capture both growth rate and yield effects (Methods, **Extended Data Fig. 1**). We conducted all experiments in biological (i.e. different overnight cultures) and technical (i.e. replicated wells in the same plate) duplicates, achieving high replicate correlations (average Pearson correlation 0.84-0.89 for biological (**Extended Data Fig. 2a-b**) and 0.91 for technical replicates (**Extended Data Fig. 2c**). Single time-point OD- and area under the growth-curve (AUC)-based fitness values led to very similar results, with the former being more robust (**Extended Data Figs. 2d-e**). Similarly, estimated single-drug fitness values mirrored experimentally-measured values and were further used since they were derived from more measurements (**Extended Data Fig. 2f-g**, Methods). From the 4 x 4 concentration matrices of fitness values, we calculated interaction scores using the Bliss interaction model^19^ (Methods, **Extended Data Fig. 1**). For each pairwise combination, a single effect size value was derived from the distribution of interaction scores, consisting of at least 72 interaction scores including all replicates of individual concentration combinations for each drug pair. The first and third quartile values of this distribution were taken as effect size values for synergies and antagonisms, respectively, with negative values corresponding to synergies and positive values to antagonisms (Methods, **Extended Data Fig. 1**, **Supplementary Table 2**)^1^. All the data is available for browsing in a user-friendly interface (https://apps.embl.de/combact/).

**Figure 2.**
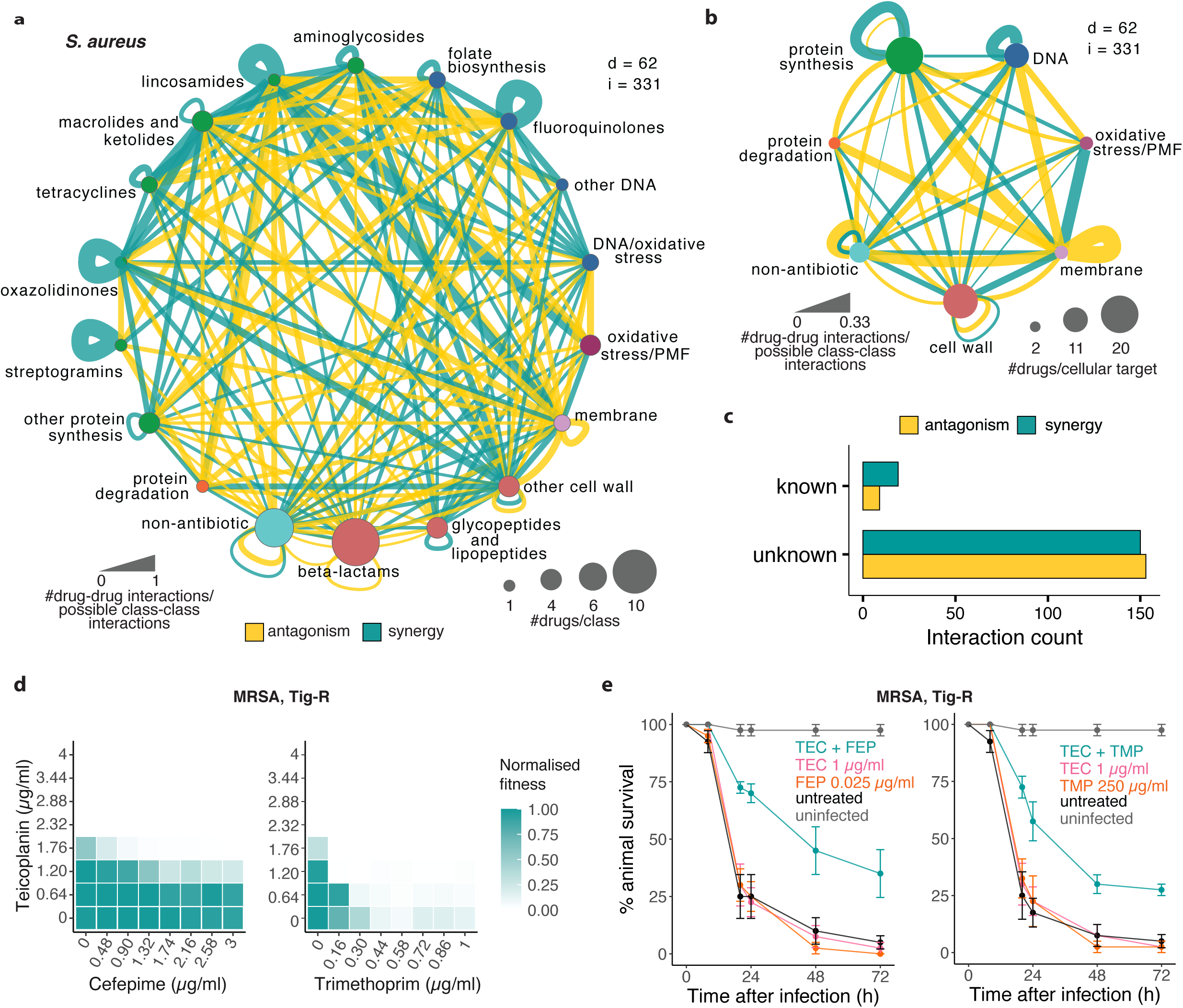
Previously uncharacterized synergies in *S. aureus* are active against clinical isolates and in infection models. **a-b**, Drug interaction networks in *S. aureus*, with drugs grouped according to their class (**a**) or targeted cellular process (**b**). Unique interactions across both strains tested are considered (i = 331). Edge thickness represents the proportion of interactions for each node pair, considering all possible interactions given the number of drugs in each node. Nodes depict the drug class (**a**) or the targeted cellular process (**b**), and size is proportional to the number of drugs in the represented class/process. Only interacting drugs are considered (d = 62). Synergies, antagonisms and nodes are coloured according to Fig. 1. **c**, Novel and previously reported interactions detected in *S. aureus.* Interactions are considered known if reported in any *S. aureus* strain (**Supplementary Table 4**). **d-e,** Novel synergies are effective against MRSA clinical isolates *in vitro* (**d**) and *in vivo* in the *G. mellonella* infection model (**e**). Teicoplanin (TEC) synergies with cefepime (FEP) and trimethoprim (TMP) were validated against a tigecycline-resistant MRSA clinical isolate (**Supplementary Table 1**) in 8 x 8 broth microdilution checkerboards (**d**) and in *G. mellonella* infection model (**e**). For checkerboards the median fitness (OD_595nm_ at 7.5h normalised by no-drug controls) across two biological replicates is shown (**Extended Data Fig. 7, Supplementary File**). For *G. mellonella* experiments, larvae were infected with the same MRSA isolate and treated with single drugs or combinations. The percentage of surviving larvae after treatment and in the untreated controls was monitored over time. Uninfected and untreated (vehicle only) controls are shown. Drugs were tested in combination at the same concentration indicated for each drug. Data points represent the mean and error bars indicate standard error (n = 10 for each condition, three independent experiments).

To calibrate hit scoring, as well as to assess the high-throughput screen data quality, we benchmarked the screen data against a validation set of 161 combinations (2% of screened combinations), equally representing the four strains probed. These combinations were tested in the same growth conditions as our high-throughput screen, but over a highly-resolved dose space (8 x 8 matrix) of equally-spaced concentration dilution gradients (Methods, **Extended Data Fig. 3a-b, Supplementary Table 2**). The precision-recall curves were comparable to the previous Gram-negative screen^1^, with highest precision (0.87) and recall (0.68) observed for a threshold on absolute effect size of > 0.1 and on Benjamini-Hochberg adjusted p-value of < 0.05 (resampling procedure with 10,000 repetitions for each combination tested, comparison with resampled Bliss scores using Wilcoxon rank-sum test in each iteration) (Methods, **Extended Data Fig. 3c**). We were able to slightly increase the recall (0.72) by relaxing the effect size thresholds for interactions found in both *S. aureus* strains, using within-species conservation as an additional parameter to confirm interactions^1^ (Methods, **Extended Data Fig. 3c-d, Supplementary Table 2**).

**Figure 3.**
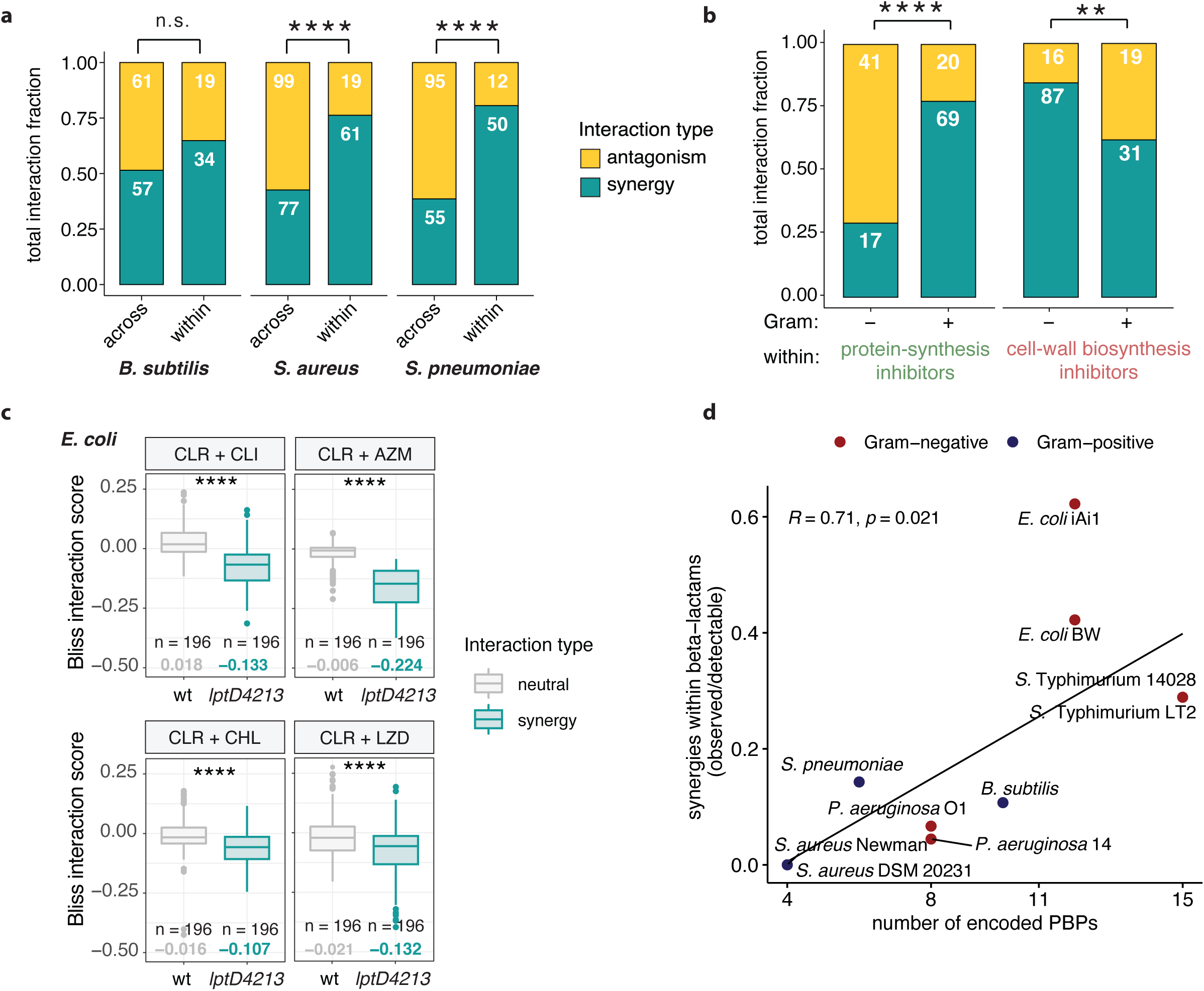
Distinct synergies between drugs targeting the same cellular process across Gram-positive and -negative species. **a**, Drugs targeting the same biological process often interact synergistically, whereas antagonisms are prevalent between drugs targeting different processes (*S. aureus*: p = 9.9e-07, χ^2^ test; *S. pneumoniae*: p = 4.7e-08, χ^2^ test). Non-antibiotic drugs (n = 8) are excluded from this analysis, as their targeted processes are heterogeneous or unknown. **b**, Gram-positive species exhibit frequent synergistic interactions between protein-synthesis inhibitors, whereas cell-wall biosynthesis inhibitors predominantly synergize in Gram-negative species. Prevalence of interactions between protein-synthesis inhibitors and between cell-wall biosynthesis inhibitors in Gram-negative and Gram-positive species (for protein synthesis inhibitors: p=1.8e-08; for cell-wall biosynthesis inhibitors: p=0.0038, χ^2^ test). **c**, Protein-synthesis inhibitors can also synergize in Gram-negative species when the drug permeability bottleneck is abolished. Synergistic combinations in Gram-positive species were tested in 8 x 8 broth microdilution checkerboards in wild-type *E. coli* and in the OM-defective *E.coli lptD4213* strain^37^. Interaction score distributions for each combination are significantly different between the two strains. Interactions were assigned with the same criteria used in the screen, with synergies corresponding to distributions with first quartile < -0.1. The first quartile value is shown in all cases. CLR, clarithromycin; CLI, clindamycin; AZM, azithromycin; LZD, linezolid; CHL, chloramphenicol (CLR + CLI: p = 2.2e-16; CLR + AZM: p = 5.3e-13; CLR + CHL: p = 2.2e-16; CLR + LZD: p = 4.4e-08, Wilcoxon test). **d**, Differences in beta-lactam synergy prevalence between Gram-negative and Gram-positive species are related to differences in drug target redundancy, that is the Penicillin Binding Proteins (PBPs) they encode in their genomes. Pearson correlation between number of PBPs (**Supplementary Table 5**) and the frequency of synergies between beta-lactams for each strain tested.

### Drug interactions are rare and species-specific

Antagonisms and synergies were detected to be equally prevalent across the three species, accounting for ∼12% of all combinations tested (**Fig. 1b**). This interaction rate, corrected by our ability to observe synergies or antagonisms according to the fitness space probed for each combination (Methods), is lower and less skewed towards antagonisms as compared to Gram-negative species (*E. coli*, *S.* Typhimurium, *P. aeruginosa*)^1^. This could be due to technical (drug or strain selection biases) or biological (Gram-positive bacteria have a lower drug permeability bottleneck than Gram-negative bacteria and hence less interactions will be dependent on depend on intracellular drug availability^1^) reasons.

Species-specificity of drug interactions has long been assumed^20^, and recently systematically demonstrated for Gram-negative species, with 30% of detected interactions shared between at least two of the three species tested, and 5% conserved in all three species (*E. coli*, *S.* Typhimurium, *P. aeruginosa*)^1^. In Gram-positive species we observed an even lower conservation rate (**Fig. 1c**), with only 81 out of 725 unique interactions (11.2%) conserved in at least two species (**Fig. 1d**). 29 interactions were conserved in all three species (4%) (**Supplementary Table 2**). We reasoned that the lower interspecies conservation in our screen could be driven by the strain and species selection in the two screens, as for Gram-negative species two closely-related enterobacteria, *E. coli* and *S.* Typhimurium, exhibited the highest overlap of interactions^1^. However, the interaction conservation rate of either of these two species with *P. aeruginosa* is similar to the cross-species conservation rates we detected for Gram-positive bacteria. Indeed, when we compared the interaction conservation rate and genome sequence percentage identity (based on 40 universal single-copy marker genes^21^), the two were significantly correlated (Methods, **Fig.1e**).

Synergies were shown to be more conserved in Gram-negative bacteria^1^, but this trend was non-significant in Gram-positive species, even after removing non-antibiotic drugs for which intracellular targets and their conservation are unknown (**Extended Data Fig. 4a)**. Conserved synergies with Gram-positive species were mostly driven by drugs targeting the same essential and highly conserved cellular processes, such as DNA biosynthesis and translation (**Fig. 1d, Extended Data Fig. 4b)**. Some of these interactions, such as the synergy between macrolides and tetracyclines or between quinolones of different generations, have been observed before in Gram-negative species^1, 22^, pointing towards conserved relationships between the targets of these compounds. Similarly, the broad antagonism between drugs targeting DNA and protein synthesis (**Fig. 1d**) is conserved in Gram-negative bacteria, and is due to the alleviation of protein-DNA imbalance after treatment with any of the two antibiotics alone^23^. Overall, we detected 52 synergies and 66 antagonisms shared across the Gram-positive/-negative divide (**Extended Data Fig. 4c-e, Supplementary Table 3**).

**Figure 4.**
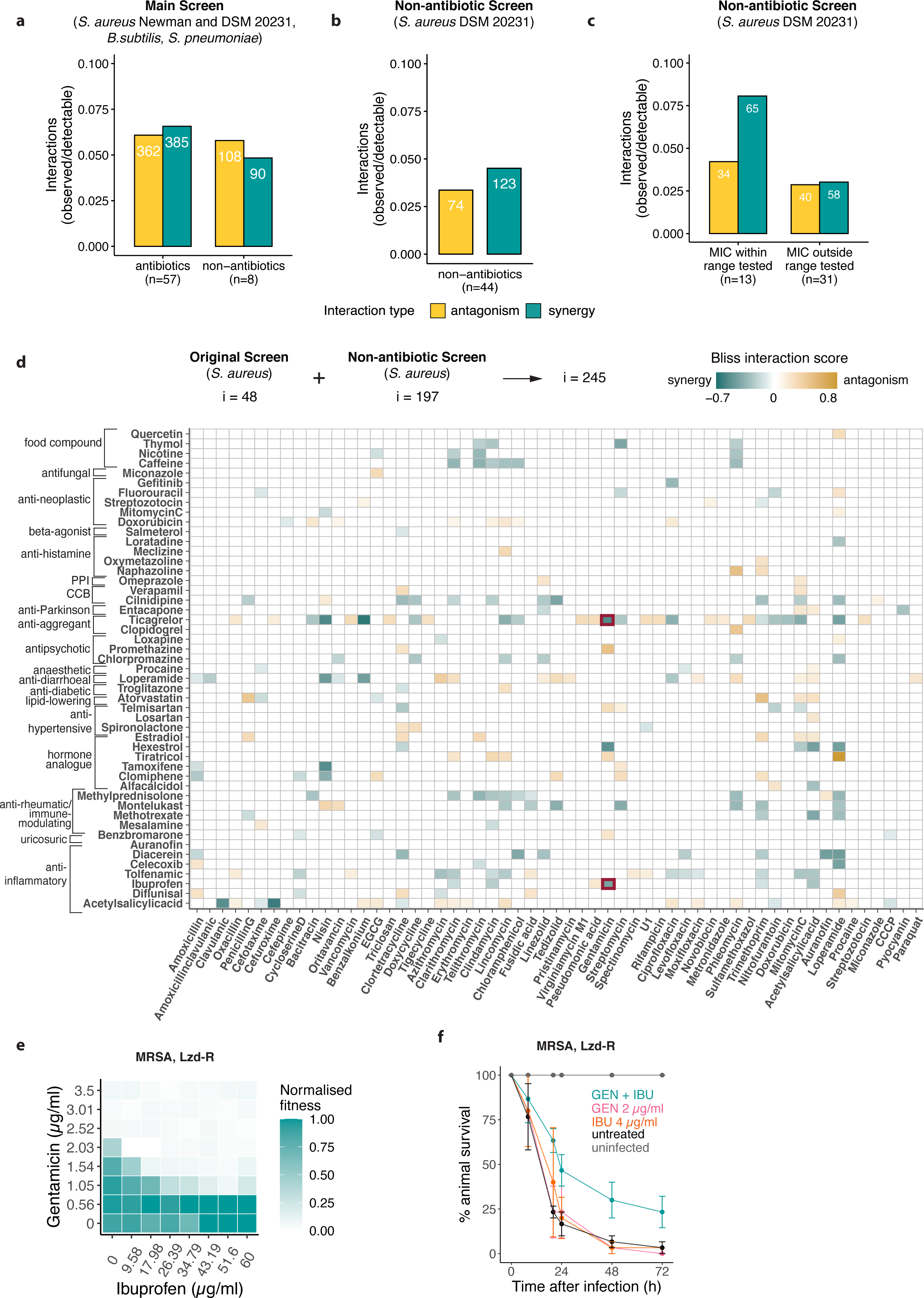
Prevalent interactions between non-antibiotic drugs and antibiotics in *S. aureus*. **a**, Interactions of non-antibiotic drugs between themselves and with antibiotics are as common as interactions between two antibiotics. This motivated us to expand the non-antibiotic panel tested. Synergy and antagonism frequencies are calculated as in Fig. 1b. **b**, Interactions between non-antibiotics and antibiotics in the extended non-antibiotic screen in *S. aureus* DSM 20231. 44 additional non-antibiotic drugs were screened in combination with 62 drugs belonging to the original drug panel, using the same experimental setup and the same data analysis pipeline, in *S. aureus* DSM 20231 (Methods). Synergy and antagonism frequencies are calculated as in Figure 1b. **c**, Non-antibiotics with antibacterial activity, for which the MIC was among tested concentrations (n = 13), engage in more interactions than non-antibiotics for which an MIC does not exist or was not within the tested concentration range (n = 31). **d**, All interactions between non-antibiotic and antibiotics detected in *S. aureus* DSM 20231 in the original (i = 87) and in the extended (i = 197) non-antibiotic screen. **e-f**, The nonsteroidal anti-inflammatory ibuprofen synergizes with gentamicin in MRSA clinical isolates *in vitro* (**e**) and in *in vivo* infection models (**f**). Gentamicin synergizes with ibuprofen and ticagrelor in an MRSA-clinical isolate with additional resistance to linezolid (**Supplementary Table 1**) in 8 x 8 broth microdilution checkerboards (**e**) and in *G. mellonella* infection model (**f**). Controls and results are obtained and represented as in Fig. 2d-e. GEN, gentamicin; IBU, ibuprofen.

### Numerous previously unknown drug synergies for *S. aureus*

We built separate interaction networks for each of the three species tested, and grouped drugs according to their class or cellular target (**Fig. 2a-b; Extended Data Fig. 5**). Although individual drug-drug interactions were only rarely conserved (**Fig. 1c**), interactions between drug classes or targeted processes were more coherent in all three species. This functional concordance became even clearer when comparing drugs based on all their interactions with other drugs. Interaction-based clustering better recapitulated drug functional classes (**Extended Data Fig. 6a**, Methods) than clustering based on chemical structures (**Extended Data Fig. 6b**, Methods), suggesting that drug interactions capture more information on drug mode of action than their chemical features.

**Figure 5.**
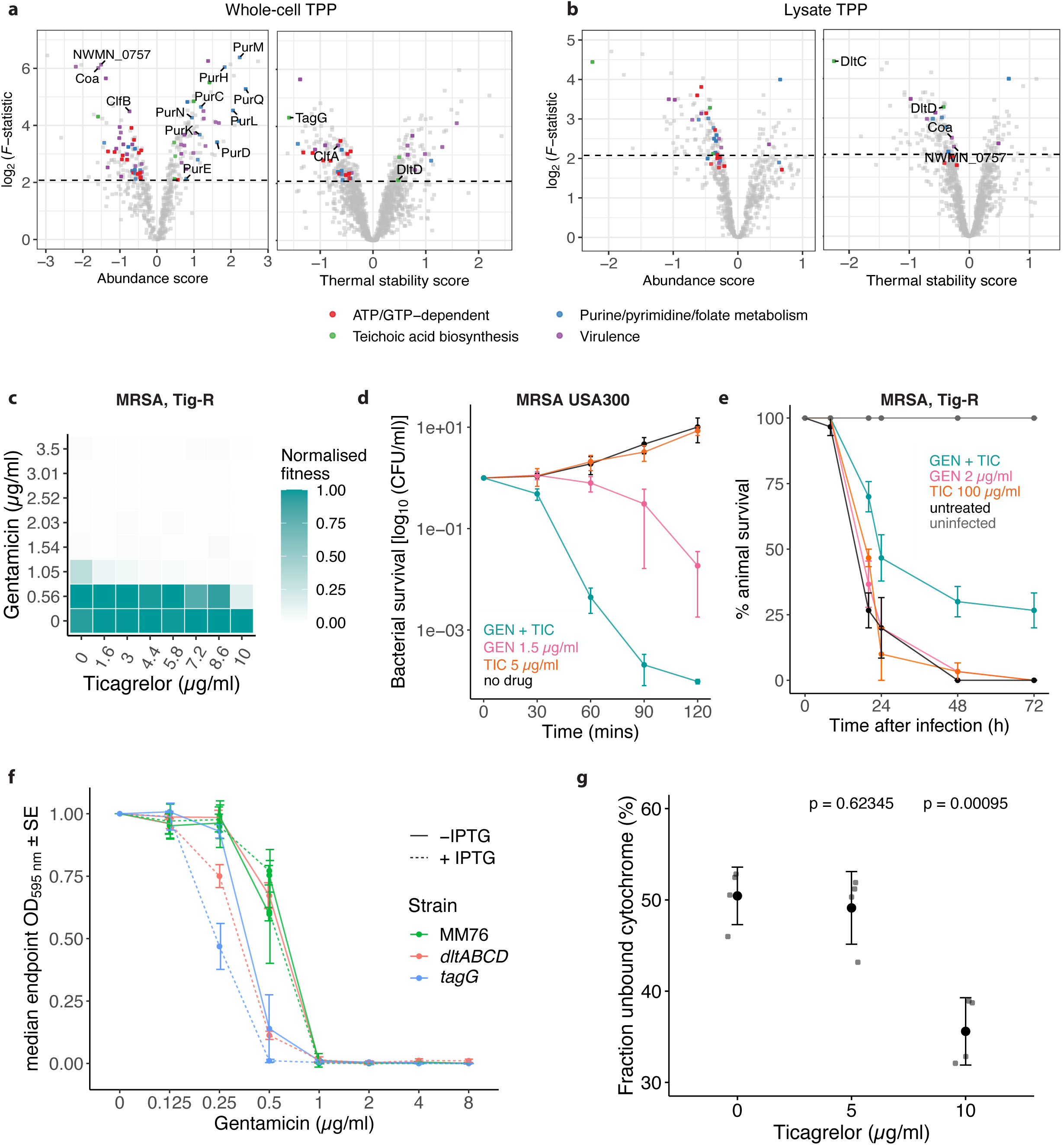
The antiplatelet ticagrelor affects *S. aureus* metabolism and synergizes with cationic antibiotics by altering *S. aureus* surface charge. **a-b**, Volcano plots highlighting abundance or stability hits in whole-cell (**a**) and lysate 2D-TPP data (**b**). The x-axis represents the effect size of protein abundance or stability change^86^ (Methods), and the y-axis corresponds to the statistical significance (log_2_(*F*-statistic)). For visualisation purposes, when the F-statistic was 0, it was transformed to 1. **c-e**, Ticagrelor synergizes with gentamicin *in vitro* at growth inhibition (**c**, **Extended Data Fig. 11a, Supplementary File**) and killing level (**d**, mean across four biological replicates; error bars represent standard error, drugs tested in combination at same concentration indicated for each drug), and *in vivo* (**e**) against an MRSA isolate resistant to tigecycline (**Supplemental Table 1**). Results for **c** and **e** are obtained and represented as described in Fig. 2d-e. GEN, gentamicin; TIC, ticagrelor. **f**, Growth (endpoint OD_595nm_, corresponding to the beginning of stationary phase for the control strain MM76, Methods, **Supplementary File**) measured in the presence of serial two-fold dilutions of gentamicin, normalised by no-drug controls, in the *S. aureus* IPTG-inducible knockdown mutants *dltABCD* and *tagG* and their control strain MM76 (Methods, **Supplementary Table 1**), in presence or absence of 500 µM IPTG to induce maximal knockdown of the gene targeted (median across four biological replicates; error bars represent standard error). All strains are grown in presence of 5 µg/ml erythromycin and 10 µg/ml chloramphenicol to maintain the CRISPRi plasmids^72^ (Methods). For all controls and full growth curves see Supplementary File. **g**, *S. aureus* Newman surface charge changes upon exposure to ticagrelor. The fraction of positively charged unbound cytochrome C is measured after incubation of drug treated and untreated samples (n = 6, mean and standard error are shown, Methods). For all controls and cytochrome c standard curve see **Extended Data Fig. 12g**.

Since *S. aureus* is the most relevant Gram-positive species in respect of AMR-attributable deaths^3^, we systematically screened literature for reported drug interactions in this species. Out of 331 unique interactions detected across the two *S. aureus* strains in our study, we found only 31 to have been previously reported (**Fig. 2c**). 55 further interactions have been reported in other bacterial species **(Supplementary Table 4**). Even when excluding those, our dataset revealed 127 novel synergies for *S. aureus* (and 118 antagonisms), a third of which (n = 39) was conserved in both strains. This confirms that the combinatorial space is a largely unexplored reservoir for improving antimicrobial efficacy.

Known interactions include many conserved synergies between drugs with the same targets (**Fig. 1d**), such as synergies between DNA-biosynthesis inhibitors, protein-synthesis inhibitors, and cell-wall targeting antibiotics (**Fig. 2a-b**). Among these latter, we confirmed the known synergy between cefepime and teicoplanin^24, 25^, and we validated it against several MRSA (Methicillin-Resistant *S. aureus*) clinical isolates, including a strain resistant to the last-resort antibiotic tigecycline (**Fig. 2d, Extended Data Fig. 7a**). When we infected larvae of the greater wax moth *Galleria mellonella* with this MRSA strain, the combination protected the animals from succumbing to the infection in contrast to single drug treatments (**Fig. 2e**), confirming that the synergies are also effective in vivo.

Synergies between cell-wall targeting drugs and translation inhibitors are cornerstones of anti-infective therapy against Gram-positive bacteria^26–29^. We could recapitulate some of these synergies, but not the traditionally used combinations of beta-lactams and aminoglycosides. This is in line with previously reported concerns on the general effectiveness of such type of combinations, whose interaction outcome seems to be strikingly strain-specific^30–33^. Other cell-wall targeting drugs may hold some unexplored potential: for instance, fosfomycin strongly synergized with a diverse range of protein-synthesis inhibitors (**Supplementary Table 4**), and could present an underexploited therapeutic resource against *S. aureus* (see Discussion).

Among the 300 previously unknown interactions we detected, 19 out of 23 tested were confirmed in extended 8 x 8 checkerboard benchmarking assays (9 of which in both *S. aureus* strains) (**Supplementary Table 2**). Interestingly, adjuvants, like clavulanic acid, or antibiotics used in clinics only in fixed-concentration combinations (trimethoprim and sulphonamides), exhibited a number of synergies with other drugs, unveiling a so-far unexplored space for new combinations. We validated the efficacy of one of these combinations (teicoplanin-trimethoprim) against several MRSA clinical isolates in vitro (**Fig. 2d, Extended Data Fig. 7b**) and in vivo in a *G. mellonella* infection model (**Fig. 2e**).

#### Fundamental differences for target-specific synergies between Gram-positive and Gram-negative species

Drugs belonging to the same class or targeting the same cellular process exhibited mainly synergistic interactions in all three species (**Fig. 2a-b, Extended Data Fig. 5, 8a**). Indeed, synergies between drugs targeting the same process were significantly enriched (**Fig. 3a**), in agreement with previous data on Gram-negative bacteria^1^. Targeting different facets of the same cellular process can bypass the inbuilt redundancy and robustness of biological processes^34^. Importantly, the targeted cellular processes that were more prone to synergies were distinct when comparing Gram-positive and Gram-negative species (**Extended Data Fig. 8a-b**). Synergies between protein-synthesis inhibitors were specifically prevalent in Gram-positive species, whereas Gram-negative species were dominated by synergies between cell-wall inhibitors (**Fig. 3b, Extended Data Fig. 8c-f**). Since the drugs between the two screens largely overlapped, and their targets are conserved in bacteria, we decided to further investigate the underlying reason for this clear difference.

Protein-synthesis inhibitors are mostly used against Gram-positive bacteria, as they often cannot cross the outer membrane (OM) of Gram-negative bacteria. We reasoned that in Gram-positive species, with no such permeability bottleneck, these drugs could synergize at their target level – the ribosome, as shown before by combinations of genetic perturbations of translation^35^. By contrast, in Gram-negative bacteria the OM permeability bottleneck likely masks such synergistic interactions and enriches for antagonisms, which are often due to a decrease in drug intracellular concentration(s)^1^. We confirmed this hypothesis by using the OM-defective *E.coli* mutant *lptD4213*, which is hyperpermeable to hydrophobic antibiotics and detergents^36, 37^. Many of the interactions between macrolides and different classes of protein synthesis inhibitors became synergistic in this *E. coli* mutant background (**Fig. 3c, Extended Data Fig. 9**), demonstrating that drug uptake bottlenecks can change antibiotic interactions.

While fosfomycin- and bacitracin-based interactions are mostly conserved within Gram-positive species and across the Gram-positive/-negative divide, interactions within beta-lactams are radically different between Gram-positive and -negative species, with synergies being rare in the former (**Extended Data Fig. 8e-f**, **Supplementary Table 3**). Beta-lactams have different affinities to the various penicillin-binding proteins (PBPs) present in bacteria^38^. Interestingly, the number and type of PBPs are largely different across bacterial species^38, 39^, so we hypothesized that this redundancy (number of PBP paralogues) was driving the observed difference. Indeed, the number of synergies in each strain tested correlated with the number of PBPs reported in each species **(Fig. 3d**). The higher the number of PBPs, the higher the probability that combining beta-lactams with different affinities to the various PBPs will lead to a synergistic bypassing of the redundancy. While further studies are needed, we hypothesize that this target redundancy drives the synergies between beta-lactam antibiotics, and that the difference we observed here between Gram-positive and -negative species likely depends on the number of PBPs in the species tested (**Supplementary Table 5**).

Altogether, these results support the concept that drug interactions mirror key properties of cellular networks, such as their functional modularity and redundancy, and reflect fundamental differences in cellular architecture and physiology across the Gram-positive/-negative divide.

### An underestimated reservoir of interactions between non-antibiotics and antibiotics

Our drug interaction screen included eight non-antibiotic drugs, which exhibited a similar interaction frequency (11%) as antibiotics (13%) (**Fig. 4a**). This motivated us to expand the panel of non-antibiotic drugs tested, and to explore the range of synergies and antagonisms antibiotics exhibit with commonly used non-antibiotic medications in *S. aureus.* We selected 44 drugs to include pharmaceuticals that can be co-administered with antibiotics in *S. aureus* infections or non-antibiotics with previously reported antibacterial activity against *S. aureus* (**Supplementary Table 6**). Altogether, we covered 19 therapeutic classes (**Fig. 4d**, **Supplementary Table 6**), testing each drug in a range of three concentrations and against the panel of 62 drugs of the initial screen (2728 drug-drug interactions, 4 x 4 dose matrix) in *S. aureus* DSM 20231. Concentrations were selected to fall within therapeutic plasma concentrations^40^, except for drugs with possible topical use, where higher concentrations were used. Interactions were scored and benchmarked as for the main screen (Methods, **Supplementary Table 6, Extended Data Fig. 10a-d).**

We confidently detected 197 interactions in the extended screen (**Fig. 4b**, **Supplementary Table 6**), an interaction frequency that was lower (7.8%) than the one of the initial screen or the set of eight non-antibiotic drugs included therein (**Fig. 4a**). Since all eight non-antibiotic drugs included in the main screen were selected because they had reported antibacterial activity, we reasoned that this could account for their higher interaction rate. Indeed, for those drugs that had antibacterial activity on their own, the interaction frequency was double (12% as compared to 5.9%) (**Fig. 4c**). For all non-antibiotics tested in this work (n = 52), we detected 140 synergies and 105 antagonisms mainly with antibiotics (**Fig. 4d**). A small number of interactions (22 synergies and 23 antagonisms) were found between two non-antibiotics. Synergies offer a so-far unexploited potential for drug repurposing, whereas antagonisms expose risks of decreasing the efficacy of antimicrobial treatments.

The therapeutic classes that exhibited the highest number of interactions were anti-inflammatory drugs (n = 7, of which four NSAIDs) and hormone analogues (n = 6) (**Fig. 4d, Extended Data Fig. 10e, Supplementary Table 6**), whereas in terms of antibiotics, protein-synthesis inhibitors dominated the interactions (**Fig. 4d, Extended Data Fig. 10f, Supplementary Table 6**). Interestingly, selective estrogen-receptor modulators (SERMs), such as the two triphenylethylene compounds tamoxifene and clomifene, shared their synergies with cell-wall acting drugs and their antagonism with streptomycin. Hormone analogues engaged in several synergies (n = 11) and antagonisms (n = 17), suggesting an understudied impact that such commonly-used drugs, and potentially their natural counterparts, may have on the efficacy of antibacterial therapies^14, 41^. For the anti-inflammatory drugs, only four interactions with acetylsalicylic acid were previously known: its synergy with cefuroxime^42^ and its antagonisms with ciprofloxacin, oxacillin and azithromycin^43–45^. We validated the synergy between ibuprofen and gentamicin also against MRSA clinical isolates, including a strain resistant to linezolid (a last-resort antibiotic for MRSA), in vitro and in vivo in a *G. mellonella* infection model (**Fig. 4e-f, Extended Data Fig. 11**).

### The antiaggregant ticagrelor has a broad impact on *S. aureus* physiology that accounts for its promiscuous interactions with antibiotics

Ticagrelor, a purine analogue antiaggregant acting on the adenosine P2Y_12_ receptor^46^, had the highest number of interactions (n = 27) among the 44 non-antibiotics tested (**Fig. 4d**). Ticagrelor has been shown to improve clinical outcomes in patients with pneumonia and sepsis caused by Gram-positive bacteria^47, 48^. This effect has been supported by different degrees of evidence that ticagrelor activates platelets upon systemic infection^48^, protects them from *S. aureus* toxin-mediated damage^49^, modulates their antibacterial properties^50^, and exerts a direct bactericidal activity on *S. aureus* at very high concentrations^51^. However, the mode of action of ticagrelor on *S. aureus* and its interactions with other drugs have remained largely uncharacterised.

To gain insights into the mode of action and interaction of ticagrelor, we used two-dimensional thermal proteome profiling (2D-TPP)^52–54^ in both lysate and whole cell samples to investigate the direct and indirect effects of the drug, respectively (Methods). We observed a destabilisation of a number of ATP- and GTP-binding enzymes and transporters in both the whole cell and the lysate (**Fig. 5a-b, Extended Data Fig. 12a, Supplementary Table 7**), and the induction of many purine biosynthesis enzymes (PurC, PurD, PurE, PurF, PurH, PurK, PurL, PurM, PurN, PurQ) in live cells (**Fig 5a, Extended Data Fig. 12a-b**). This is in agreement with ticagrelor being a purine analogue. Furthermore, the MIC of ticagrelor increased upon supplementation of defined media with adenosine, inosine or their combination (**Extended Data Fig. 12c**), suggesting that ticagrelor interferes with purine metabolism in *S. aureus*.

As mentioned above, the clinically observed effects of ticagrelor during *S. aureus* infection have not been linked so far to a direct effect of ticagrelor on *S. aureus* virulence. We discovered a pervasive impact of ticagrelor on *S. aureus* virulence determinants and regulators, many of which were down-regulated, and others destabilized (**Fig 5a, Extended Data Fig. 12a**, **Supplementary Table 7**). In particular, we observed destabilisation in lysate and downregulation in whole-cell samples of key clotting factors secreted by *S. aureus* (ClfA, ClfB), the coagulase Coa and the von Willebrand-factor binding protein (vWBP) NWMN_0757^55^. These effects, evident at a clinically-relevant ticagrelor concentration^51^ offer an alternative explanation for the beneficial effect of antiaggregant therapy as an adjuvant in *S. aureus* systemic infection.

Ticagrelor exhibited a number of synergies and antagonisms with antibiotics in MSSA (Methicillin-sensitive *S. aureus*; **Fig. 4d**). Interestingly, it broadly sensitized MSSA and MRSA to both cationic peptides (nisin; **Extended Data Fig. 12d**) and antibiotics (aminoglycosides, such as gentamicin; **Fig. 5c, Extended Data Fig. 11**). This potentiation effect of aminoglycosides occurred at low ticagrelor concentrations, and was also evident at the killing level (**Fig. 5d)** and in vivo, during infection of *G. mellonella* (**Fig. 5e**). Since aminoglycosides need energy to cross the membrane in most bacteria^56^, we wondered whether ticagrelor acted at that level, for example by modulating the cell surface charge and increasing aminoglycoside uptake. Consistent with this hypothesis, two proteins involved in the lipoteichoic acid (LTA) D-alanylation^57^, DltC and DltD, were destabilized in the TPP lysate data, and TagG, a subunit of the cell wall teichoic acid (WTA) translocase, was destabilised in the whole cell sample (**Fig 5a-b, Extended Data Fig. 12e**). Disruption of teichoic acids, and specifically, inactivation of *dltA*, *dltB* and *dltC* have been shown to sensitize *S. aureus* to cationic compounds because of an increase in the net negative charge of *S. aureus* surface^58–60^. We observed a decrease in the MIC of the aminoglycoside gentamicin and the cationic antibiotic nisin in IPTG-inducible CRISPRi knockdown mutants of both the *dltABCD* operon and of *tagG* (Methods, **Fig. 5f, Extended Data Fig. 12f**). Ticagrelor treatment also increased the binding of positively-charged cytochrome C to intact *S. aureus* cells (**Fig. 5g, Extended Data Fig. 12g-h**). Thus, ticagrelor treatment impacts the thermal stability and presumably the activity of proteins involved in WTA flipping and LTA D-alanylation, leading to an increase in the surface net negative charge of *S. aureus*. This leads to potentiation of the uptake of cationic antibiotics, such as aminoglycosides and nisin.

## Discussion

In this study, we have systematically profiled drug combinations in three prominent Gram-positive bacterial species. We probed multiple compounds from each of the main antibiotic classes used to treat infections caused by Gram-positive pathogens, as well as neglected antibiotics, commonly used antibiotic adjuvants, and promising non-antibiotic drugs with reported antibacterial activity. In all cases combinations were tested in a dose-dependent manner and interactions were assessed in a quantitative manner. This effort unravelled a plethora of interactions, the majority of which have not been previously not reported. A number of the synergies that we discovered using lab strains were also effective against MDR clinical isolates and during infections in vivo. Overall, the data generated here can seed future experiments to mechanistically dissect key interactions or to explore their potential for clinical application. For example, some of the synergies and antagonisms identified may guide future broad-spectrum empiric treatments – i.e. when antibiotic regimens are started without knowledge of the responsible pathogen in time-sensitive contexts (e.g. sepsis). Fosfomycin synergies that are strong and conserved across the Gram-positive/-negative divide are good candidates, as fosfomycin is increasingly used in clinics^61^, but rarely in combinations. To enable further use of this rich resource, we made all data browsable in a user-friendly interface.

A previous large-scale study from our group in three Gram-negative species^1^ enabled important comparative insights across the Gram-positive/-negative divide. The confidence and depth level of these comparisons are high, since the two studies have similar experimental and data analysis design (including the drugs tested). As in Gram-negative species, drug interactions are largely species-specific with synergies tending to be more conserved and driven by antibiotics sharing general cellular targets. Only a small number of interactions is conserved across Gram-positive and -negative species. Differences in the cell surface organization (e.g. the outer membrane posing a permeability barrier for hydrophobic compounds) or in the degree of redundancy in cell-wall building enzymes can explain some of the strong synergies observed specifically in Gram-positive or Gram-negative species.

We also attempted to leverage the adjuvant potential of approved non-antibiotic drugs by probing a large number of combinations with antibiotics (2728) in a dose-dependent manner in *S. aureus*. Although the interaction potential was lower for drugs without antibacterial activity, the room for novelty is high with the vast majority of synergies detected being previously unknown. While non-antibiotic drugs have been proposed as anti-infective adjuvants for decades^5, 6^, the *in vivo* relevance and molecular basis of their antibacterial action is known only for a few examples^6, 8, 18, 51, 62, 63^. Here we further studied the antiaggregant ticagrelor, whose repurposing as an anti-infective adjuvant for Gram-positive bacteria has been recently proposed^49, 64^. While the *in vivo* benefit of ticagrelor for systemic infections has been documented^47, 51^, we exposed here a large number of additional synergies with antibiotics in *S. aureus* (n = 13), and provided molecular insights into how ticagrelor affects *S. aureus* physiology and potentiates the activity of positively-charged antibiotics, such as aminoglycosides or nisin.

Drugs are regularly combined in clinics not only in rationally designed therapeutic schemes, but also extemporarily in poly-treated patients^14^. Although interactions at the host pharmacokinetic level are routinely avoided, it is assumed that interactions at the bacterial level would not affect overall anti-infective efficacy. In addition to synergies, we detected an equally large number of antagonisms between commonly administered non-antibiotic drugs and antibiotics. Such antagonisms could decrease the efficacy of the antibiotic treatment and increase the probability of resistance emergence for the antibiotic. Overall, it is important to start assessing drug interactions not only at the level of growth inhibition, but also at the level of killing and ultimately clearing of the infection, as the outcome of interactions may differ^11^.

It has also been recently proposed that the attenuation of antibiotic efficacy (antagonism) can be used to decrease the spectrum and collateral damage of antibiotics to commensal bacteria^65^. In our screen, loperamide had the highest number of interactions with antibiotics. Although its potential use as adjuvant for specific antibiotics and its mode-of-action are known^6^, we could detect an additional broad antagonism with macrolides. Loperamide and macrolides are often co-administered for travellers’ diarrhoea^62^, which is caused by Gram-negative enteric pathogens. It is tempting to speculate that part of the beneficial effect of the combination could also lie in the protection of Gram-positive commensal gut species from macrolides.

In summary, we present a systematic and quantitative account of drug interactions in key Gram-positive species, discovering a number of potent synergies that are effective against clinical MDR isolates, and providing insights into the underlying mechanisms of some of the observed interactions. In an era where novel antibiotic development faces technical and economic hurdles, and new antimicrobial strategies are urgently needed, systematic drug interaction profiling can offer possible solutions to treat bacterial infections. Extending the systematic testing of drug interactions to more bacteria and to other types of non-antibiotic drugs will improve our understanding of drug interaction conservation and mechanisms, and potentially inform tailored treatments towards pathogens.

## Materials and methods

### Strains and growth conditions

All strains used in this study are listed in **Supplementary Table 1**. *B. subtilis* subsp *subtilis* 168^66^ was kindly provided by Carol A. Gross, all MRSA clinical isolates by Stephan Göttig, *S. pneumoniae* D39V^67^ by Jan-Willem Veening, and *S. aureus* USA300 by Daniel Lopez. *Staphylococcus aureus* subsp *aureus* Newman^68^ was purchased from NCTC (NCTC 8178). and DSM 20231^69^ (ATCC 12600 ^T^, NCTC 8532) from DSMZ, Braunschweig, Germany.

For all experiments and unless otherwise specified, *S. aureus* strains were grown in Tryptic Soy Broth (TSB, ref. 22092 by Merck-Millipore), *B. subtilis* was grown in LB Lennox, and *S. pneumoniae* was grown in CY medium, adapted from^70^. All species were grown at 37°C, with vigorous shaking (850 rpm), except for *S. pneumoniae*, which was grown without shaking. The ticagrelor purine supplementation experiments in *S. aureus* Newman were conducted in SSM9PR defined medium supplemented with 1% glucose^71^.

### Inducible knockdown strain construction

A two-plasmid CRISPR interference system was used to knock down gene expression of selected genes in *S. aureus* Newman^72^. In these strains, *dcas* is expressed from an IPTG-inducible promoter on plasmid pLOW, while sgRNAs are expressed from a constitute promoter on a plasmid derived from pCG248. The sgRNA-target sequences were TGTCTAACAGCAATGCTTTG for *dltABCD* and AAACCATAATTTGCATAACA for *tagG*, and ATAGAGGATAGAATGGCGCC for the non-target control MM76 (**Supplementary Table 1**).

### MIC determination

MICs were tested in all strains for the main screen (**Supplementary Table 1**). Drugs were two-fold serially diluted in 11 concentrations, and 32 no-drug control wells were included in each plate. Experiments were conducted in flat, clear-bottom 384 well plates (ref. 781271 by Greiner BioOne), with a total volume of 30 µl for *S. aureus* and *B. subtilis* and 55 µl for *S. pneumoniae*. Volumes were optimised for each strain to achieve good dynamic range for growth and minimize risk of cross-contamination between wells. Plates were inoculated with a starting optical density at 595 nm (OD_595nm_) of 0.01 from an overnight culture. All liquid handling was performed using a Biomek FX liquid handler (Beckman Coulter). Plates were sealed with breathable membranes and incubated at 37°C. OD_595nm_ was measured every 30 minutes for 14 hours. The MIC was considered as the first concentration at which growth was inhibited. Experiments were conducted in biological duplicates. For the adjuvant screen, antibiotics were tested in *S. aureus* DSM 20231 at the same concentrations as for the main screen. Non-antibiotics concentrations were selected to fall within therapeutic plasma concentrations^40^(**Supplementary Table 6)**.

### High-throughput screen of drug combinations

Sixty-two drugs, hereafter designated as recipients, were arrayed in flat, clear-bottom 384-well plates in three two-fold serial dilutions and two technical replicates – up to two recipient drugs were removed from the data of the different strains due to quality control reasons. Concentrations were selected according to MICs, with the highest concentration corresponding to the MIC, and the intermediate and lowest concentration corresponding to half and a quarter of MIC, respectively (**Supplementary Table 1**). Plates were kept frozen and defrosted upon each experimental run, when the same 62 drugs and in the same three concentrations were added as donor drugs (one drug at one concentration for each recipient plate). A few drugs were screened only as donors: the combinations co-amoxiclav and cotrimoxazole in *B. subtilis* and *S. aureus* DSM 20231; co-amoxiclav, clavulanic acid, pseudomonic acid and cefuroxime in *S. aureus* Newman. All donor drugs were tested in two biological replicates. Control wells were included in each plate (six no-drug wells, three plain medium wells and three wells containing only the donor drug). After the addition of donor drugs, plates were inoculated with cells. Handling, inoculation, growth conditions, plate incubation and OD_595nm_ measurements were performed as for the MIC determination.

For the adjuvant screen, 44 non-antibiotic drugs (**Supplementary Table 6**) were tested against the same 62 recipient drugs of the main screen for *S. aureus* DSM 20231.

### Screen benchmarking and 8 x 8 checkerboard assays

Combinations were tested in the same experimental conditions as for the screen, but at higher concentration resolution. Drugs were diluted in eight concentrations spanning linearly-space gradients to assemble 8 x 8 checkerboards for each combination tested – highest concentration used can be found in **Supplementary Table 2**. All experiments were conducted in at least two technical and two biological replicates. Data was analysed with the same pipeline as for the screen (Methods).

### Data analysis

Data analysis was adapted from^1^. Growth curves were processed as described in **Extended Data Fig. 1**: the background was subtracted from all OD_595nm_ measurements, on a well-by-well basis using the first measurement obtained. Abnormal spikes in OD values of the first three time points occurred in *S. pneumoniae* in a small fraction of wells due to bubble formation in the medium or plate condensation. These early local peaks in OD curves were identified and replaced with the median of OD values (of corresponding time points) estimated from the wells not affected by such artefacts within the same plate. When more than one in the first four time points was affected, those wells were identified as non-monotonically-increasing OD_595nm_ values across the first four time points, and their background was estimated as the median first-time-point OD_595nm_ of artefact-free wells (monotonically increasing across the first four time points).

The time point corresponding to the transition between exponential and stationary phase in no-drug control wells was identified as the first time point at which the maximum OD_595nm_ was reached. This time point was selected according to the growth characteristics of each strain, and was kept the same across runs: 8 h for both *S. aureus* strains, 5.5 h for *B. subtilis* and 3.7 h for *S. pneumoniae*. Later time points were excluded from further analysis. The OD_595nm_ measurement at that time point was used to derive fitness measurements that captured effects both on growth rate and maximum yield. OD-based endpoints correlated well with AUC-based endpoints (Pearson correlation 0.95), and led to higher precision and recall according to the screen benchmarking (**Extended Data Fig. 2d-e**).

This value was then divided, per plate, by the robust mean^73^ of the six no-drug controls (no-drug control hereafter), obtaining three fitness measures for each drug concentration pair: *f*_1_, fitness upon exposure to drug 1; *f*_2_, fitness upon exposure to drug 2; and *f*_1,2_, fitness in presence of drug 1 + drug 2. Based on these values, further quality control was again performed, correcting fitness increase artefacts (maximum fitness was set to 1) and removing plates with poor technical replicate correlation (Pearson correlation < 0.7). *f*_1_, *f*_2_, and *f*_1,2_ were used to calculate interaction scores using the Bliss model^19^. The choice of this model over other available quantification methods was driven by the following considerations: (i) the three measurements obtained for drug dose responses are not sufficient for accurate quantification using alternative models (e.g. the Loewe model^74^) and (ii) the Bliss model can more accurately account for single drugs with no effect (such as most non-antibiotic drugs included in the screen).

Bliss (ε) scores were calculated as follows:

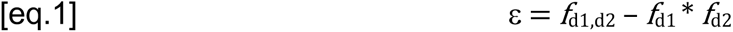

where *f*_d1d2_ corresponds to the observed fitness in the presence of the drug combination, and *f*_d1_ and *f*_d2_ correspond to the fitness in the presence of the two single drugs.

Single drug fitness for both donor and recipient drugs can also be inferred from combination fitness, by minimizing the sum of residuals squared of the Bliss independence model as follows and using the assumption that most drugs interact neutrally:

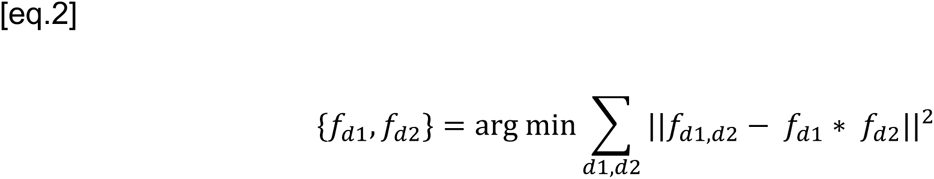

Experimentally measured and estimated fitness values were very similar for donor (**Extended Data Fig. 2f**) and recipient (**Extended Data Fig. 2g**) drugs, and we used the estimated measures since those were more robust to noise - experimental controls were limited for donor drugs (three single-drug control wells) and sometimes biased for recipient drugs, as a single problematic plate in the batch was sufficient to generate noise.

When no data was discarded upon quality control, the number of Bliss scores obtained for each combination was 72, composed of 3 x 3 (in the two-dimensional concentration space) x two technical replicates x two biological replicates x two replicates with drugs tested as donor or recipient. Hit calling was performed using a resampling procedure with 10,000 repetitions for each combination tested, where the ε distribution for each combination was compared with the resampled Bliss scores using Wilcoxon rank-sum test in each iteration^1^. Hits correspond to combinations with FDR < 0.05.

As before^1^, we coupled this significance threshold to an effect-size threshold. For each combination we defined a cumulative score using the quartiles of its distribution of ε scores. We tested the performance of different thresholds in precision and recall upon screen benchmarking, and identified |0.1| as best threshold, with precision 0.87 and recall 0.68 (**Extended Data Fig. 3d**). Accordingly, synergies were assigned if the first quartile of the ε distribution < -0.1 and antagonisms if the third quartile exceeded 0.1. We could increase the screen recall by leveraging the presence of two strains belonging to the same species in the case of *S. aureus*, as previously described for Gram-negative species^1^. We defined an additional set of hits (weak and conserved), meeting significance and effect size thresholds in one strain, but with lower effect size in the other strain. A cutoff of |0.08| allowed us to maintain the same precision and increase the recall to 0.72.

Data analysis was implemented with R v.4.1.2^75^ and RStudio v.2021.09.1^76^ and networks were created with Cytoscape v.3.8.2^77^.

### Interaction detection calculation

Interaction detection rates were calculated by dividing the number of detected interactions by the number of combinations for which interactions could be observed according to the mapped fitness space^1^. Synergies could not be observed when the expected fitness of a drug combination (defined as the product of single-drug fitness values in eq. 1) was lower than 0.1, while antagonisms could not be detected for expected combination fitness higher than 0.9 (2.1 and 3.3% of the 7986 combinations tested, respectively).

### Drug clustering

Drug-drug interaction profiles were clustered according to the cosine similarity of quartile-based Bliss interaction scores of each drug pair in each strain. Scores from all interactions were considered, regardless of their statistical significance. For the clustering based on chemical structures, drugs were clustered according to their Tanimoto similarity^78^ using 1024-bit ECFP4 fingerprints^79^.

### Phylogeny analysis

To calculate the percentage sequence identity between bacterial species, the genomes of *B. subtilis* 168, *S. aureus* Newman, *S. aureus* DSM 20231, *S. pneumoniae* D39, *E. coli* K-12, *S. enterica* serovar Typhimurium LT and *P. aeruginosa* PAO1 were downloaded from NCBI and 40 universal single-copy marker genes (MGs) were extracted using the fetchMG script^80^. The 40 MGs were selected from a previous publication for their ability to characterise prokaryotic species^21^, and they encode for ubiquitous functions like tRNA synthetases or are ribosomal proteins (EggNOG COGs: COG0012, COG0016, COG0018, COG0048, COG0049, COG0052, COG0080, COG0081, COG0085, COG0087, COG0088, COG0090, COG0091, COG0092, COG0093, COG0094, COG0096, COG0097, COG0098, COG0099, COG0100, COG0102, COG0103, COG0124, COG0172, COG0184, COG0185, COG0186, COG0197, COG0200, COG0201, COG0202, COG0215, COG0256, COG0495, COG0522, COG0525, COG0533, COG0541, COG0552). The concatenated sequences (all six genomes contained exactly 40 MGs) were used to calculate percentage nucleotide sequence identity with vsearch^81^ and to create a phylogenetic tree. To this end, a multiple sequence alignment was created with muscle v3.8.1551^82^ with default parameters. Finally, a maximum-likelihood phylogenetic tree was constructed using the online tool PhymL 3.0^83^ with default parameters. To evaluate interaction conservation, only the 46 drugs tested both in Gram-positive and Gram-negative species (**Supplementary Table 1**) were considered.

### Evaluation of drug combination therapy using the *G. mellonella* infection model

Larvae of the greater wax moth (*Galleria mellonella*) at their final instar larval stage were used for evaluation of selected drug combinations to assess their efficacy against MRSA *in vivo*. Larvae were purchased from UK Waxworms (Sheffield, UK) and Mucha Terra (Ahaus-Altstätte, Germany). Stock solutions of cefepime, gentamicin, ibuprofen, teicoplanin and trimethoprim were freshly prepared as described for the *in vitro* experiments (**Supplementary Tables 1, 6**) with the exception of ticagrelor which was dissolved in 50 mM EtOH and diluted in distilled water to the required concentration. Drug toxicity was preliminarily assessed injecting larvae with serial dilutions of single drugs and combinations. Concentrations at which no toxicity was observed were selected for further experiments. The MRSA strains were cultivated in brain heart infusion medium and harvested at an OD_600_ of 0.5. Bacteria were washed twice with PBS and adjusted to an OD_600_ which corresponded to a lethal dose of approximately 75% (LD75) of the larvae after 24 h (approximately 10^7^ CFUs). Ten larvae per condition were injected with 10 µL of the bacterial cell suspension or PBS (referred to as uninfected control) into the hemocoel via the last left proleg using Hamilton precision syringes. After one hour, 10 µL of single drugs combinations, or vehicle were injected into the last right proleg, at the following drug concentrations: teicoplanin 1 µg/ml, trimethoprim 250 µg/ml, cefepime 0.025 µg/ml, gentamicin 2 µg/ml, ibuprofen 4 µg/ml, ticagrelor 100 µg/ml. The survival of *Galleria* larvae was monitored at the indicated time points by two observers independently. Each strain–drug combination was evaluated in three independent experiments.

### Time-kill experiments

Overnight cultures of *S. aureus* USA300 were diluted 1:100 in 20 ml of TSB medium, incubated for 1h in flasks at 37°C with continuous shaking and diluted again 1:100 in 20 ml prewarmed TSB with ticagrelor (5 µg/ml), gentamicin (1.5 µg/ml), their combination or without drugs. 50 µl of serial 10-fold dilutions of cultures were plated on TSA plates every 30 minutes for 2 hours. Cell viability was determined by counting CFUs after plates were incubated overnight in four independent experiments.

### Two-dimensional thermal proteome profiling (2D-TPP)

Bacterial cells were grown overnight at 37°C in TSB and diluted 1000-fold into 50 ml of fresh medium. Cultures were grown at 37°C with shaking until OD_578_ ∼0.6. Ticagrelor at the desired concentrations (0.04, 0.16, 0.8 and 4 µg/ml) or a vehicle-treated control were added and cultures were incubated at 37°C for 10 minutes. Cells were then pelleted at 4,000 x *g* for 5 min, washed with 10 ml PBS containing the drug at the appropriate concentrations, resuspended in the same buffer to an OD_578_ of 10 and aliquoted to a PCR plate. The plate was then exposed to a temperature gradient for 3 min in a PCR machine (Agilent SureCycler 8800), followed by 3 min at room temperature. Cells were lysed with lysis buffer (final concentration: 50 μg/ml lysostaphin, 0.8% NP-40, 1x protease inhibitor (Roche), 250 U/ml benzonase and 1 mM MgCl_2_ in PBS) for 20 min, shaking at room temperature, followed by five freeze–thaw cycles. Protein aggregates were then removed by centrifuging the plate at 2,000 x *g* and filtering the supernatant at 500 x *g* through a 0.45 µm filter plate for 5 minutes at 4°C. Protein digestion, peptide labelling, and MS-based proteomics were performed as previously described^53^.

### 2D-TPP data analysis

Data were pre-processed and normalized as previously described^52^. Peptide and protein identification were performed against the *S. aureus* Newman strain Uniprot FASTA (Proteome ID: UP000006386), modified to include known contaminants and the reversed protein sequences. Data analysis was performed in R using the package TPP2D^84^ as previously described^85^. Briefly, to identify stability changes, a null model, allowing the soluble protein fraction to depend only on temperature, and an alternative model, corresponding to a sigmoidal dose-response function for each temperature step, are fitted to the data. For each protein the residual sum of squares (RSS) of the two models are compared to obtain an F-statistic. FDR control is performed with a bootstrap procedure as previously described^85^. The abundance or thermal stability effect size was calculated for each protein as following:

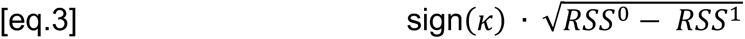

where κ is the slope of the dose-response model fitted across temperatures and drug concentrations and RSS^0^ and RSS^1^ correspond to the residual sum of squares of the null (pEC50 linearly scaling with temperature) and alternative model, respectively^86^.

### KEGG enrichment

*S. aureus* Newman proteome was annotated using KEGG^87^ (release 100.0, October 1). Proteins with missing KEGG annotation were preliminarily removed. Fisher’s exact test was then used to test the enrichment of input protein sets (hits corresponding to FDR < 0.05) against the background (all detected proteins) for each term. The p-values were corrected for multiple testing using the Benjamini-Hochberg procedure. The analysis was performed in R using the packages KEGGREST^88^, EnrichmentBrowser^89^ and clusterProfiler^90^.

### Ticagrelor MIC upon purine depletion and supplementation

Ticagrelor (ref. SML2482, Sigma-Aldrich) MIC was measured upon purine supplementation in *S. aureus* Newman as described above in SSM9PR defined medium supplemented with 1% glucose^71^, in flat, clear-bottom 384-well plates with a final volume of 30 µl. Adenine and inosine were added at 20 and 100 µg/ml or in combination, both at 100 µg/ml. Experiments were conducted in four biological replicates. A single time-point OD_595nm_ at the transition between exponential and stationary phase (13.5 h) was used to derive dose-response curves, after normalisation to the respective no-drug control for each condition.

### Gentamicin and nisin MIC measurements in *dltABCD* and *tag*G knockdown mutants

For gentamicin and nisin MIC measurements, *dltABCD* and *tagG* IPTG-inducible knockdown mutants (Methods, **Supplementary Table 1**) were grown in two-fold dilutions of nisin and gentamicin, in presence of erythromycin (5 µg/ml) and chloramphenicol (10µg/ml) for plasmid maintenance. IPTG (500 µM) was used to achieve maximal dCas9 expression and thereby, knockdown of the gene targeted. The parent *S. aureus* Newman and the control strain MM76 (containing the two vectors with dCas9 and a non-targeting sgRNA) were included in all experiments, and experiments were conducted in four biological replicates in 384-well plates. For each plate, we identified the time point where the control strain MM76 (in presence of erythromycin, chloramphenicol and IPTG at the above-mentioned concentrations) reached plateau, defined as the first time point before no increase was detected in log_10_(OD_595nm_) values of two consecutive time points. This time point was then used for all wells to derive dose-response curves, after normalisation to the respective no-drug control for each strain and biological replicate. Full growth curves annotated with the time point used for the dose-response curves and dose-response curves with all controls are included in the **Supplementary File.**

### Determination of cell surface charge

The cytochrome c binding assay was conducted as previously described^91^. Briefly, overnight cultures of *S. aureus* Newman were diluted 1:1000 in 20 ml of TSB medium, and grown in flasks at 37°C with continuous shaking until they reached OD_578nm_ ∼ 0.45. Samples were then incubated in the same conditions with or without 10 and 5 µg/ml ticagrelor for 20 minutes. Samples were centrifuged at 10000 g for 15 minutes at room temperature, washed twice with 20 mM MOPS buffer (pH 7) and concentrated to reach a final A_578_ of 10 in a 96 well-plate (ref. 4483481, Applied Biosystems™) containing cytochrome c (0.25 mg/ml, ref. 101467, MP Bio) or MOPS buffer (**Fig. 5g**). The plate was incubated in the dark at room temperature for 10 minutes. The cell pellets were collected, and the amount of cytochrome c in the supernatant was determined spectrophotometrically at an OD_410nm_. Two-fold dilutions of cytochrome c in the same plate, starting from 256 µg/ml, were used to obtain a standard curve onto which a linear model was fitted to calculate cytochrome c concentrations in the other wells. Results are expressed as unbound cytochrome c fraction in the supernatants. Experiments were conducted in four biological replicates.

## Data and code availability

Drug combination data and the computational pipeline are available on Github: https://github.com/vladchimescu/comBact.

An interactive interface to navigate the screen data is available (https://apps.embl.de/combact/).

The mass spectrometry proteomics data have been deposited to the ProteomeXchange Consortium via the PRIDE partner repository with the dataset identifier PXD036188.

## Acknowledgements

We thank the EMBL Proteomics Core Facility for supporting the 2D-TPP experiments; the Typas lab and in particular Vallo Varik for helpful discussions; Carol Gross, Jan-Willem Veening and Daniel Lopez for strains; Heike Brötz-Oesterhelt for providing the compound ADEP4^92^; Stephan A. Sieber for providing the compound U1^93^; Sara Riedel-Christ for her help with *G. mellonella* infection experiments; Zhian Salehian for his support in the construction of the *S. aureus* inducible knockdown mutants. This work was supported by EMBL and the JPIAMR grant COMBINATORIALS to A.T., JPIAMR grant DISPRUPT to Mo.K. and A.T. Mi.K. was supported by a fellowship from Swedish Research Council, A.Me. & K.M. by a fellowship from the EMBL Interdisciplinary Postdoc (EI3POD) programme (MSCA COFUND; 664726).

## Author contributions

E.C. & A.T. conceived and designed the study. O.P.K., Mo.K., G.Z., M.M.S., S.G., W.H. and A.T. supervised the study. E.C. & A.T. designed the experiments: E.C. performed the MIC testing; E.C., K.I. & A.B.-N. performed the two high-throughput screens; E.C. & A.B.-N. did the benchmarking for the screens; E.C. tested all clinical isolates and performed the follow-up work with translation inhibitors in *E. coli*; E.C., Mi.K. & J.S. performed the follow-up work with ticagrelor, except for the TPP experiments which were conducted by A.Ma., E.C. and K. M., and the CRISPRi knockdown construction, which was carried out by M.T.M.; M.T. and S.G. designed and performed the *G. mellonella* infection experiments. V.K. implemented the screen data analysis pipeline and data visualization interface, with input from W.H. E.C. & V.K analyzed the data from both screens, with input from A.R.B. A.Mi. performed the phylogeny analysis. A.Ma. and E.C. analyzed the TPP data. E.C. designed figures, with inputs from V.K., Mi.K., A.Ma., A.T. E.C. and A.T. wrote the manuscript with input from all authors. All authors approved the final version.

## Competing Interest declaration

EMBL has filed a patent application on drug combinations identified in this study ("Novel combinations of antibiotics and non-antibiotic drugs effective in vivo against Gram-positive bacteria, in particular methicillin-resistant *S. aureus* (MRSA)", European patent application number EP22207154.0). E.C., S.G. and A.Ty. are listed as inventors.

## Author information

Correspondence and requests for materials should be addressed to.

## Extended data figure legends

**Extended Data Fig. 1.**
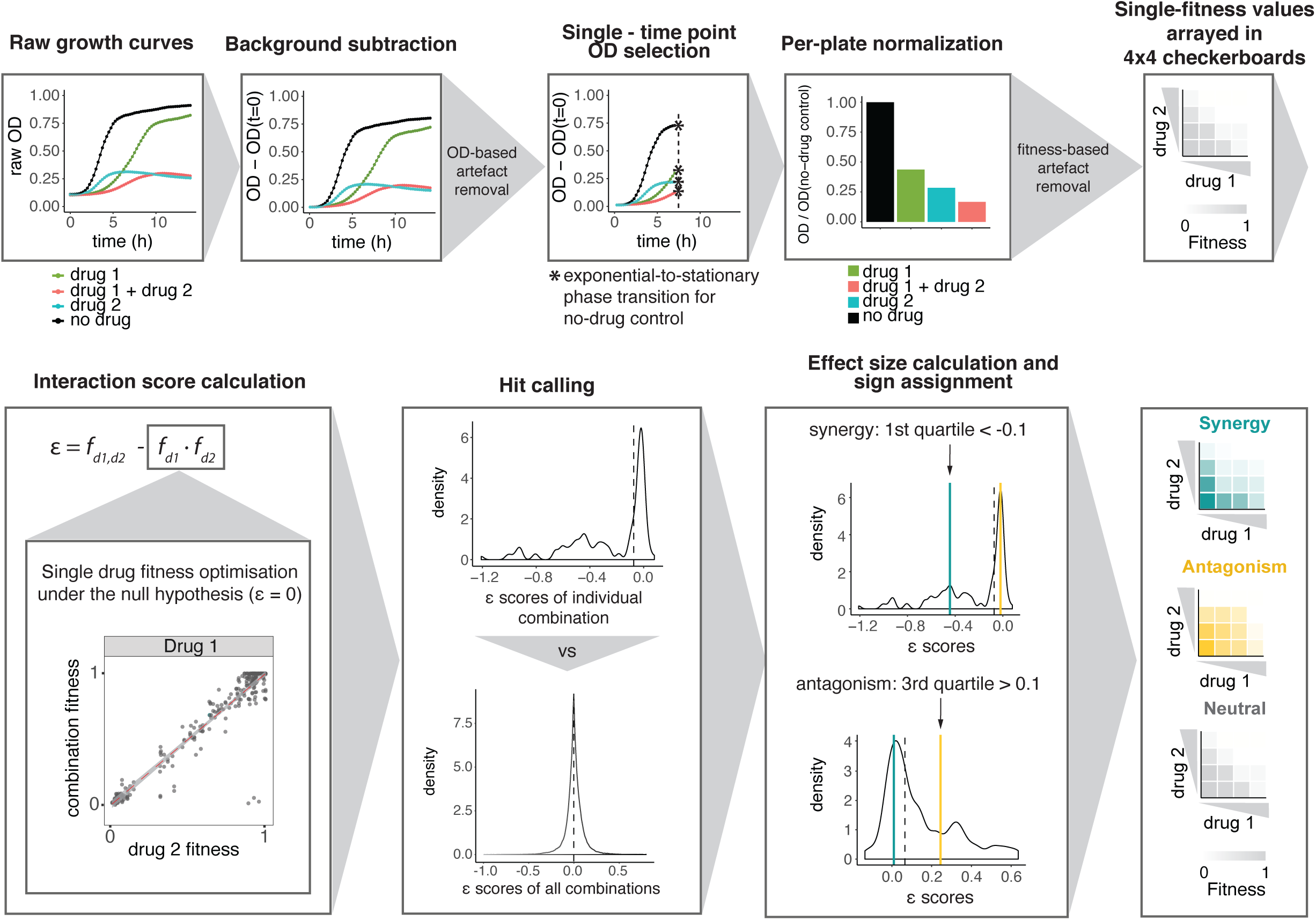
Data analysis pipeline. Raw growth curves based on measurement of OD_595nm_ over 14h were processed as depicted. Background was removed by subtracting the OD_595nm_ at the first time point (when this was not affected by artefacts) from all the following measurements (Methods). All curves within a plate were trimmed beyond the point that the no-drug controls within the plate (6 wells) entered stationary phase. The OD_595nm_ measurement at this time-point was then normalised per plate by the robust mean of the no-drug control wells (6 per plate), resulting in fitness values (Methods) that were used to obtain 4 x 4 checkerboards for each combination. Bliss (ε) scores were then calculated as follows: ε = *f*_d1,d2_ – *f*_d1_ * *f*_d2_, where *f*_d1,d2_ corresponds to the observed fitness in the presence of the drug combination, and *f*_d1_ and *f*_d2_ correspond to the fitness in the presence of each single drug. Single-drug fitness values were estimated from drug-combination fitness by minimizing the sum of residuals squared of the Bliss independence model (Methods, [eq. 2]). Interactions fulfil two criteria: (i) FDR < 0.05, after applying a resampling procedure with 10,000 repetitions of a two-sided Wilcoxon rank-sum test, to compare the ε distribution of each combination tested to the overall ε distribution; and (ii) a quartile-based effect size threshold examining the ε distribution of each combination, with synergies assigned if first quartile (green line) < -0.1 and antagonisms if third quartile > 0.1 (yellow line).

**Extended Data Fig. 2.**
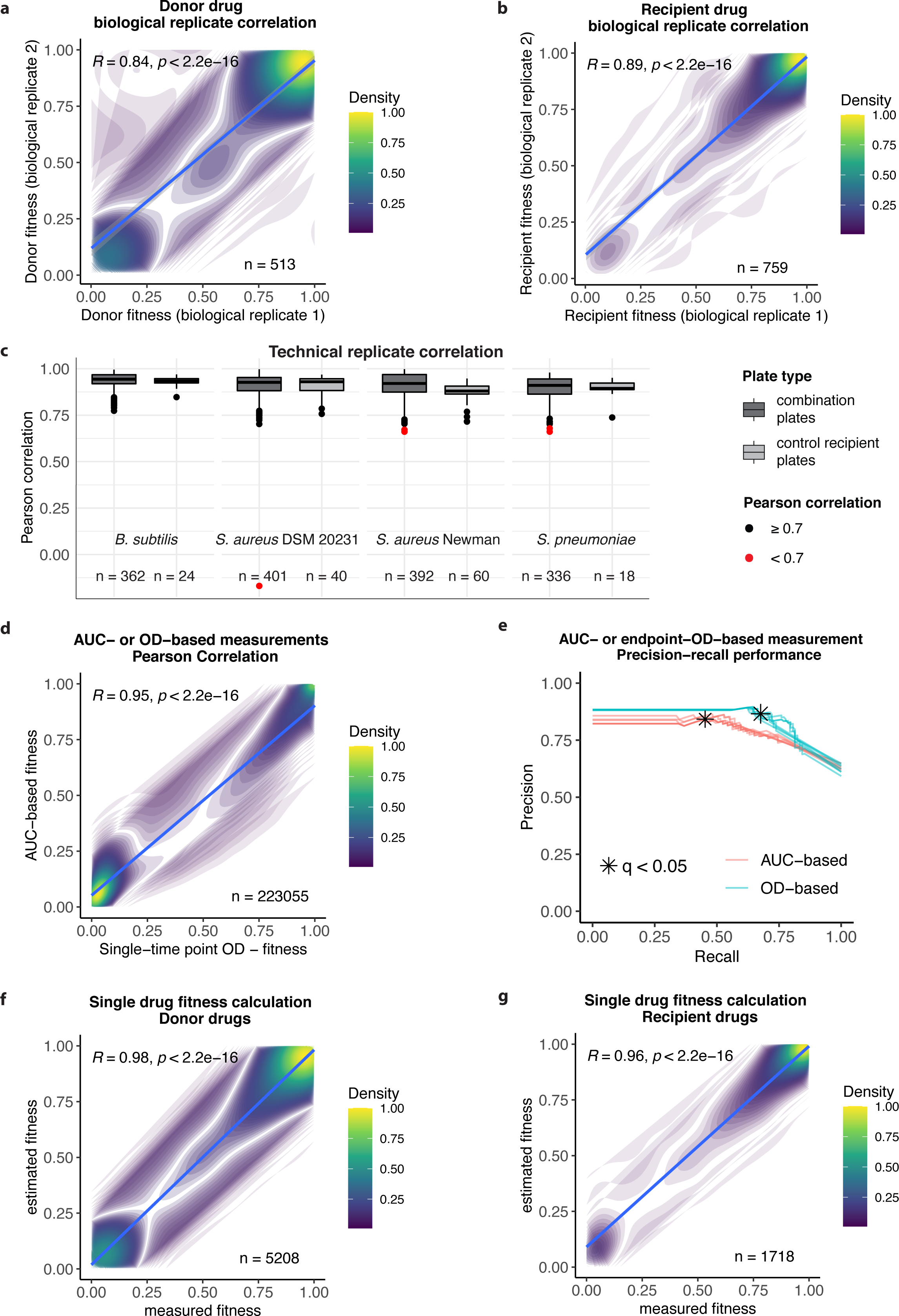
Quality control of the main interaction screen and assessment of fitness calculation methods. **a-b**, Donor (**a**) and recipient (**b**) drug fitness correlation between biological replicates. Pearson correlation is calculated between biological replicates, corresponding to different experimental runs/batches. **c**, Technical replicate correlation. Pearson correlation is calculated between replicate wells within the same plate for combination plates (where donor drugs were added) and control recipient plates (where no donor drug was added), for the four strains screened. Plates for which technical plate correlation was < 0.7 (red) were removed from the data. **d**, Endpoint-OD- and AUC-based fitness values for all strains are highly correlated (Pearson correlation, R = 0.95, p < 2.2e-16, n = 223055). **e**, Performance of endpoint OD- and AUC-based measurements in terms of precision-recall against the benchmarking set. Precision-recall curves are shown for q-value intervals increasing by 0.01. Curves highlighted correspond to the effect-size cutoff selected for the screen (interaction score = |0.1|). The selected significance cutoff (FDR < 0.05) is marked. **f-g**, Comparison between estimated and experimentally measured single-drug fitness for donor (**f**) (Pearson correlation, R = 0.96, p < 2.2e-16, n = 5208) and recipient (**g**) drugs (Pearson correlation, R = 0.98, p < 2.2e-16, n = 1718).

**Extended Data Fig. 3.**
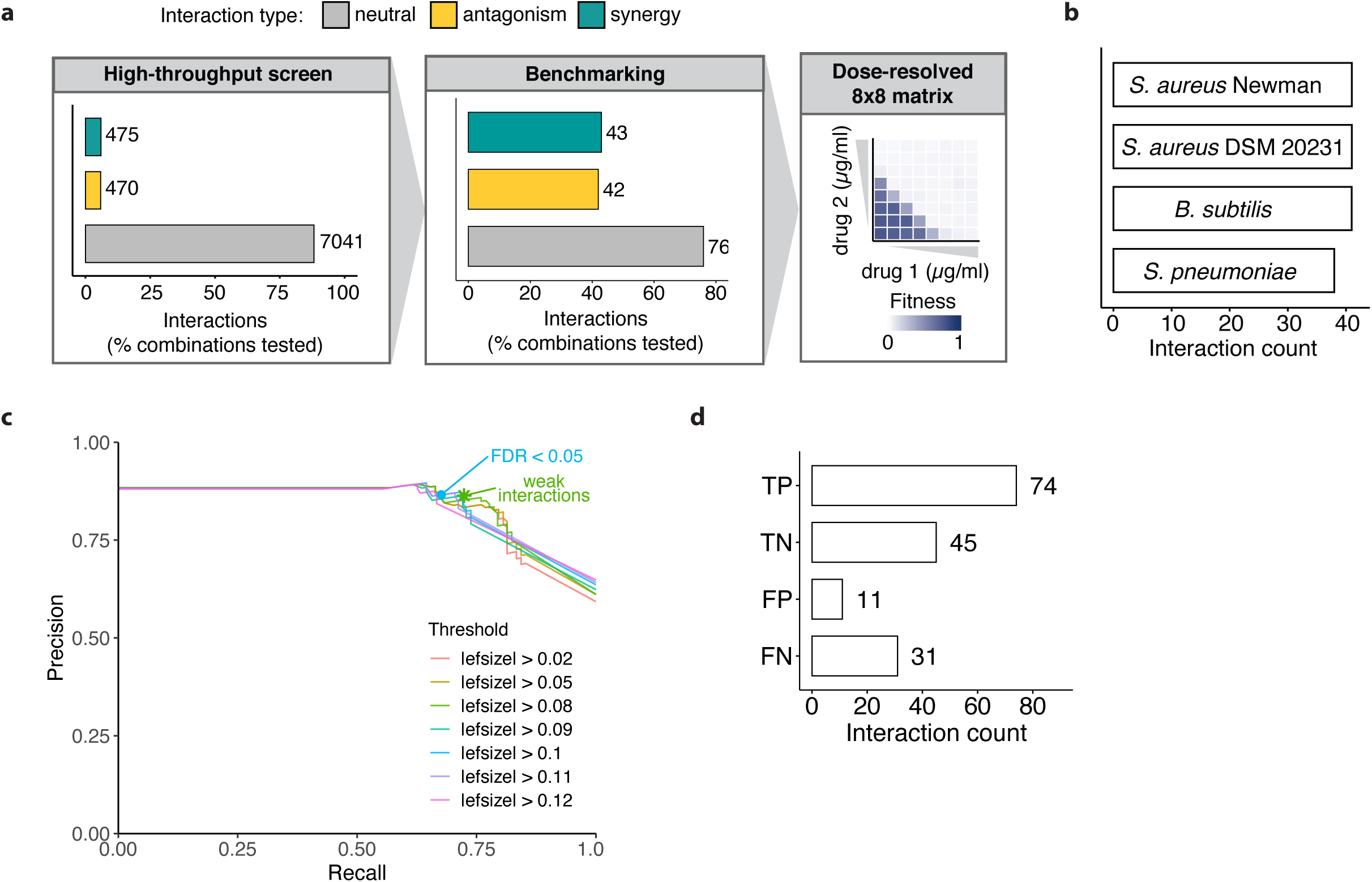
Screen benchmarking. **a**, 161 drug combinations were selected for benchmarking, including hits and neutral interactions, and tested in extended concentration checkerboards (8 x 8). Fitness values and interaction scores were calculated as in the high-throughput screen (Methods). **b**, Combinations were selected to equally represent the four strains tested. **c**, Screen precision and recall against the benchmarking set are assessed for different effect-size thresholds. Precision-recall curves are shown for FDR intervals ranging from 0 to 1, increasing by 0.005. The chosen significance value for the screen (FDR < 0.05) is highlighted for the effect-size curve (|0.1|) providing best precision and recall. The addition of weak interactions (effect-size threshold |0.08|, Methods) increases slightly the recall. |efsize|=effect size. **d**, True-positive (TP), true-negative (TN), false-positive (FP) and false-negative (FN) abundance in the benchmarking set for optimal thresholds. As for most screens, conservative cutoffs for interactions minimize FPs with a cost on the number of FNs.

**Extended Data Fig. 4.**
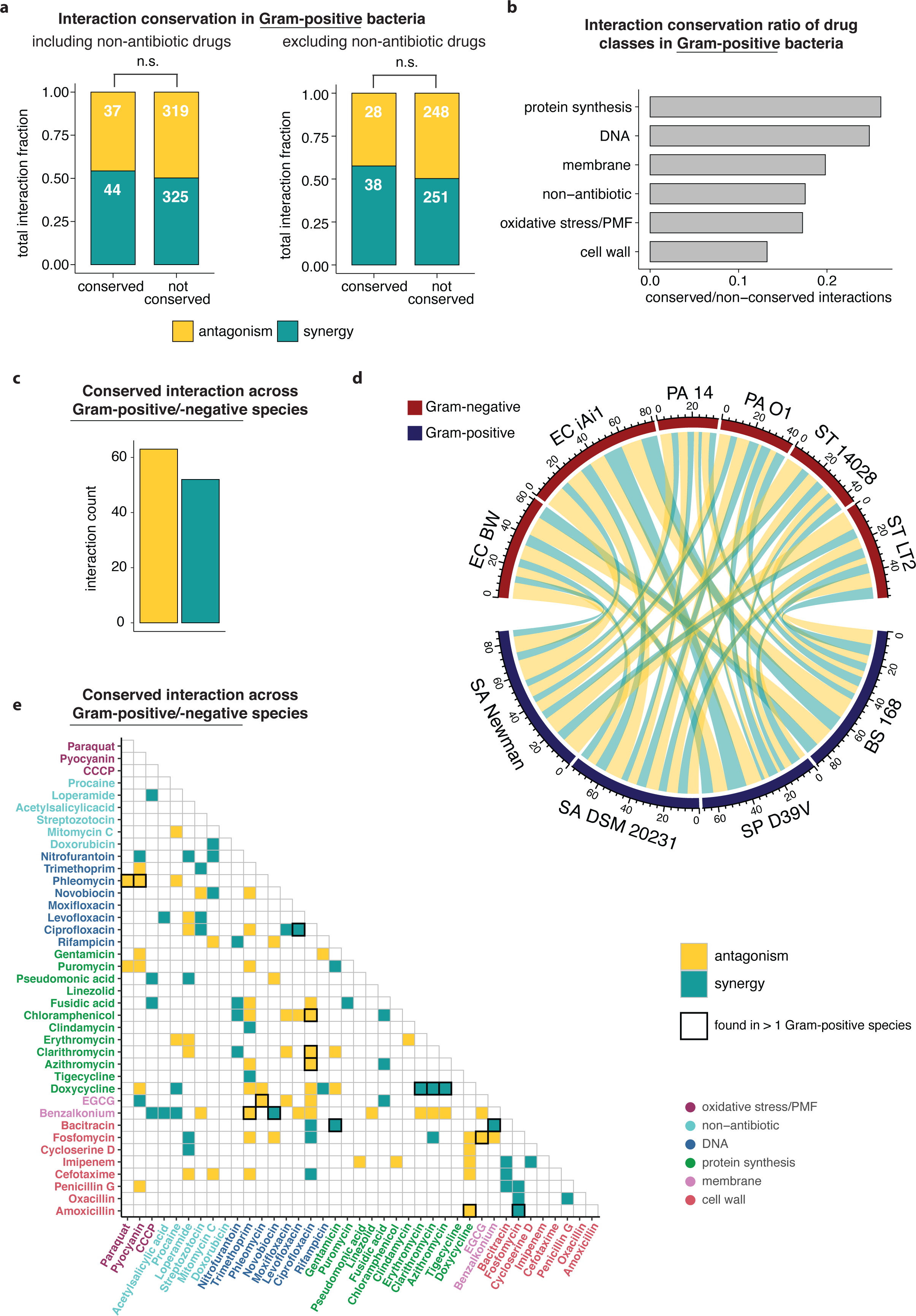
Interaction conservation within Gram-positive species and across the Gram-positive/-negative divide. **a**, There is no significant difference between synergy and antagonism prevalence among conserved and non-conserved interactions, regardless of whether non-antibiotic drugs, whose targets are multiple or unknown, are considered (p = 0.592, χ^2^ test) or not (p = 0.327, χ^2^ test). Only interactions conserved across at least two species are considered (n = 81, Fig. 1c). **b**, Drugs targeting more conserved cellular processes tend to have more conserved interactions. Interaction conservation ratio for each drug class across species is calculated as the ratio between conserved and non-conserved interactions. **c**, Synergy and antagonism abundance of unique interactions shared by at least one Gram-negative and Gram-positive strain – edges are colour-coded according to whether interaction is synergistic (green) or antagonistic (yellow). **d**, Conserved interactions across at least one Gram-negative and Gram-positive strain. EC, *E.coli*; PA, *P. aeruginosa*; ST, *S.* Typhimurium; SA, *S. aureus*; SP, *S. pneumoniae*; BS, *B. subtilis*. **e**, Heatmap of conserved interactions across Gram-positive and Gram-negative bacteria. Interactions that are also conserved across multiple Gram-positive species are highlighted.

**Extended Data Fig. 5.**
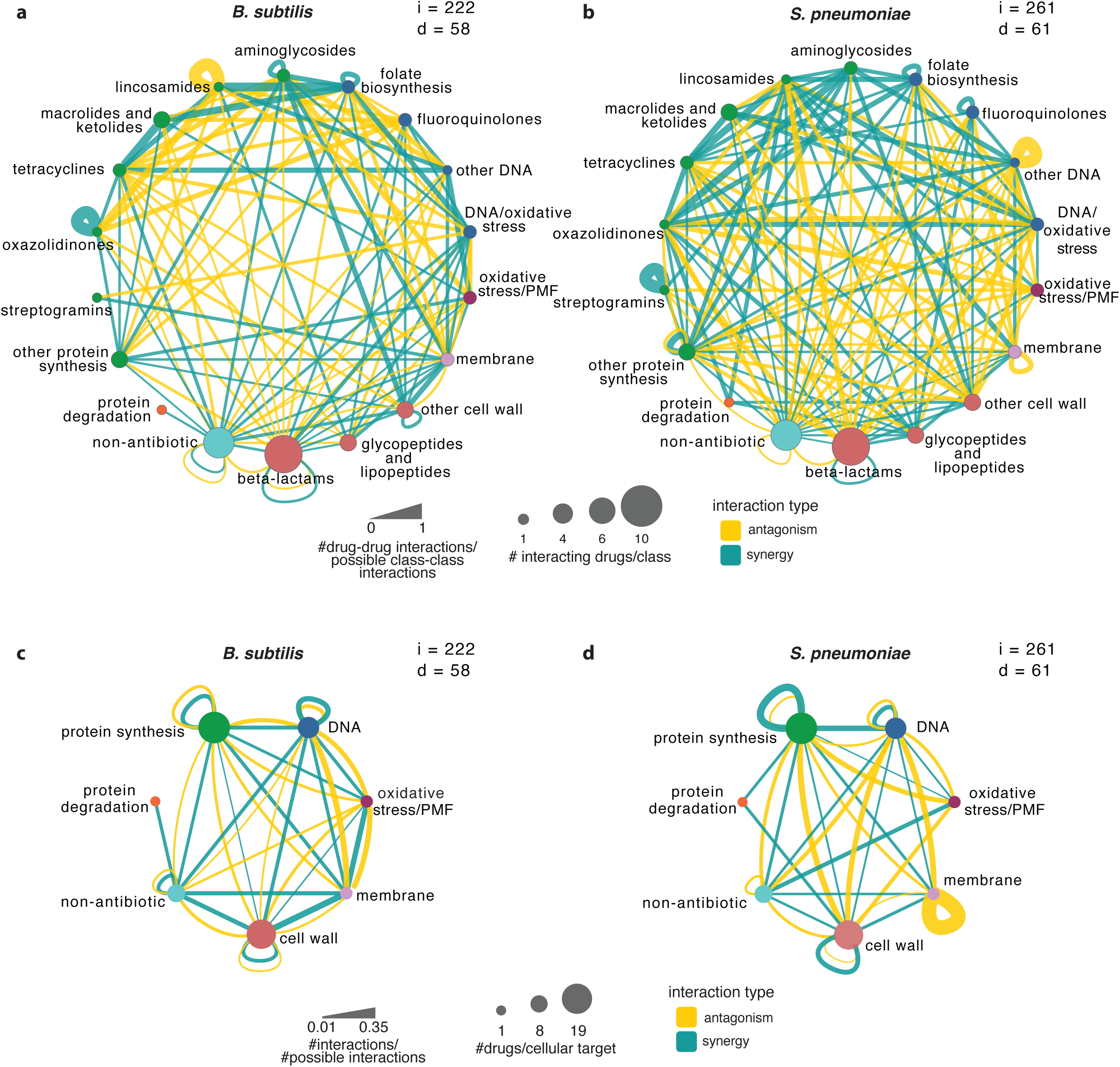
Interaction networks with drugs grouped according to their class or their targeted cellular processes for *B. subtilis* (a, c) and *S. pneumoniae* (b, d). Node size and colour and edge thickness are depicted as in Fig. 2a-b. i, number of interactions; d, drugs involved in the interactions (may differ from number of drugs tested in screen).

**Extended Data Fig. 6.**
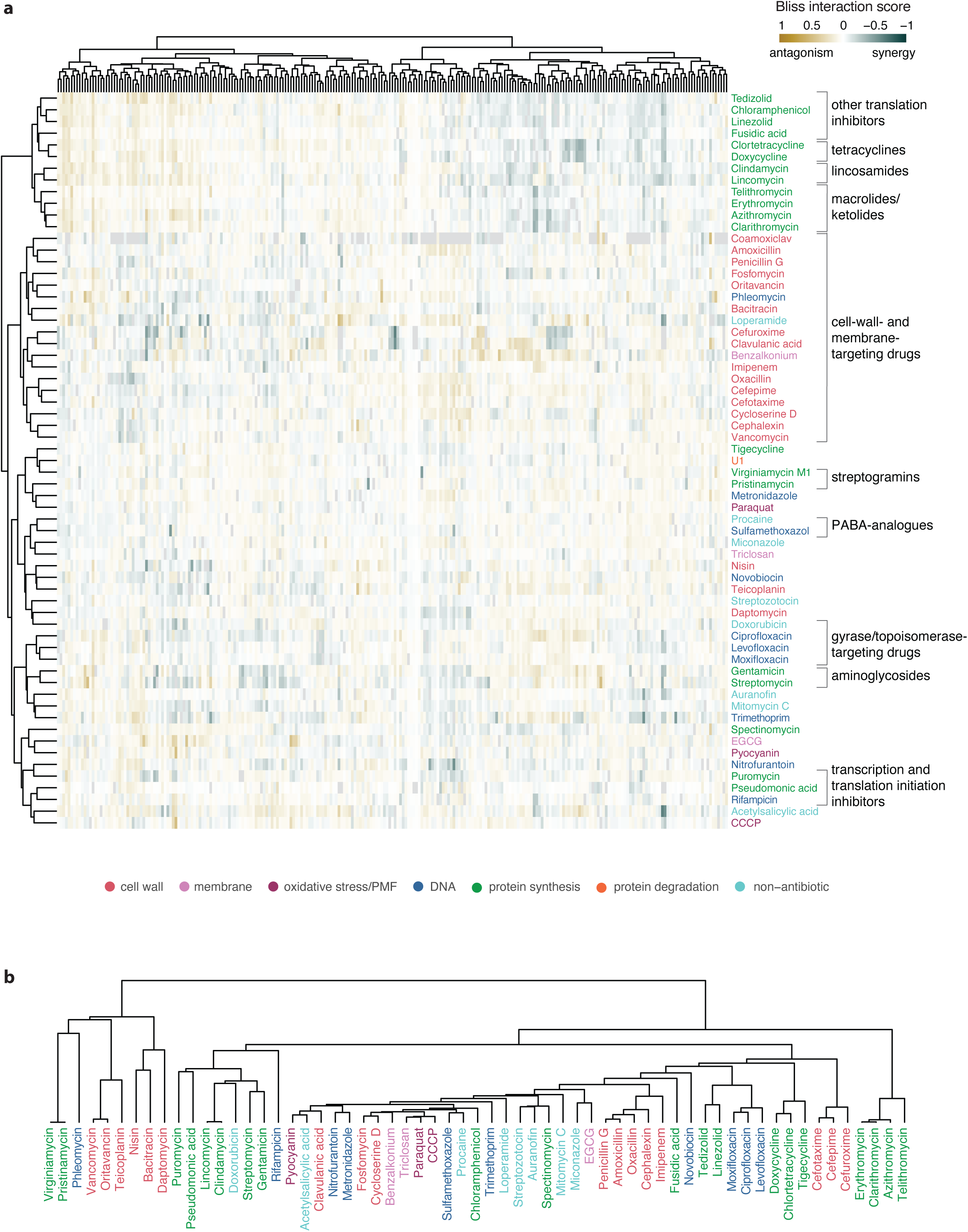
Drug interaction fingerprint recapitulates their functional and chemical classes. **a**, Drugs clustered according to their interactions with all other drugs in main screen. For each drug, quartile-based Bliss interaction scores (Methods) with all the other drugs (n = 65) in all four strains (x-axis) are considered; drug interaction fingerprints are then clustered according to their cosine similarity. All interaction cumulative scores are considered, regardless of their significance. Clusters enriched in drugs belonging to the same classes, targeting the same processes, and/or chemically similar, are highlighted. Negative, positive and neutral Bliss scores are depicted in shades of green, yellow, and in white, respectively. Combinations that were not tested in a given strain are in grey. **b**, Drug clustering according to their chemical structure similarity (Methods). In both panels, drugs are coloured according to their targeted cellular process (colour code as in Fig. 1).

**Extended Data Fig. 7.**
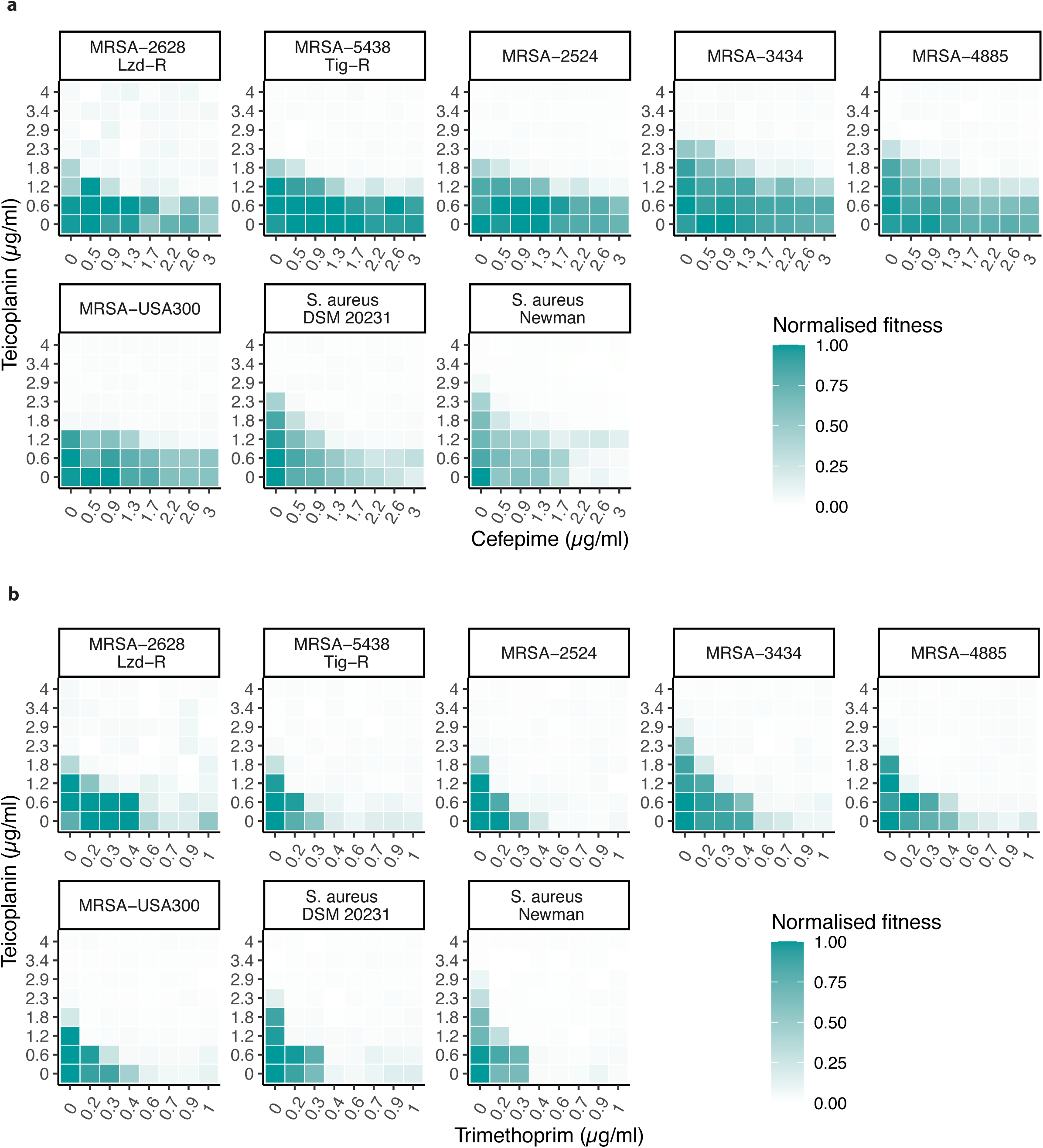
Antibiotic synergies detected in screen are effective against MRSA clinical isolates. Teicoplanin synergies with cefepime (**a**) and trimethoprim (**b**) were tested in 8 x 8 checkerboards in model MSSA (Newman and DSM 20231) and MRSA (USA300) strains, and in 5 clinical MRSA strains from different worldwide-prevalent clonal complexes, isolated from different infection sites, and bearing different multidrug resistance profiles (**Supplementary Table 1**). Results are obtained and represented as in Fig. 2d. One representative checkerboard of n = 2 (biological replicates) is shown here (for the second replicate see Supplementary File). Lzd-R, linezolid-resistant; Tig-R, tigecycline-resistant.

**Extended Data Fig. 8.**
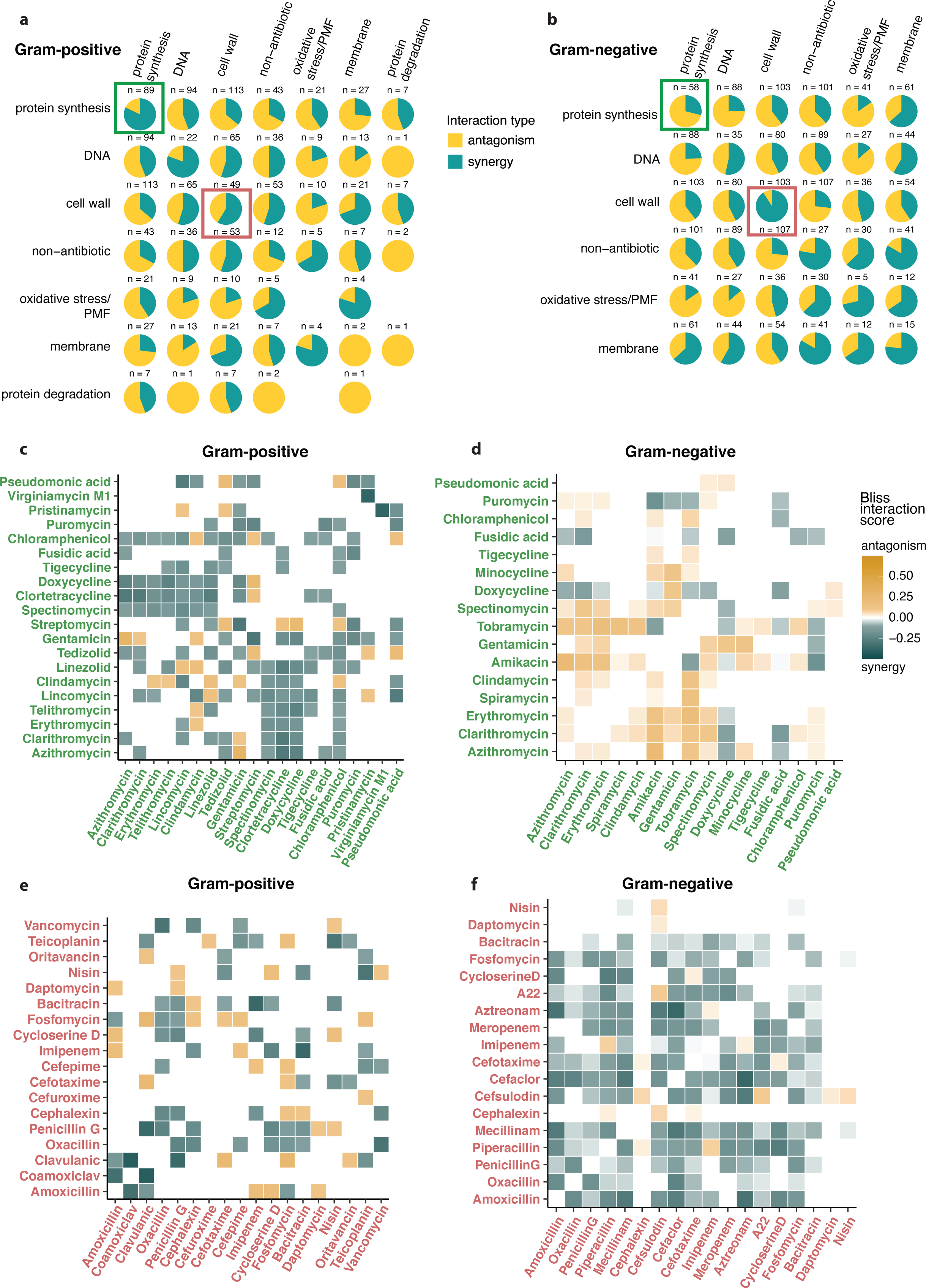
Interactions between drug functional classes in Gram-positive and Gram-negative species. **a**, Interactions between all drug classes (based on cellular target) in Gram-positive (**a**) and Gram-negative (**b**) species. The absolute count for each class-class interaction is indicated. PMF = proton-motive force. Interactions between drugs tested in all strains are considered. Interactions conserved across different strains are considered as distinct occurrences. **c-f**, Heatmaps of interactions between protein synthesis inhibitors (**c-d**) and between cell-wall biosynthesis inhibitors (**e-f**). Bliss interaction scores are averaged across strains if the same interaction is found in more than one strain. In rare cases in which opposite interactions are found in two different strains (n = 4, **c**; n = 2, **d**; n = 2, **e**; n = 8, **f**), the strongest one is shown here.

**Extended Data Fig. 9.**
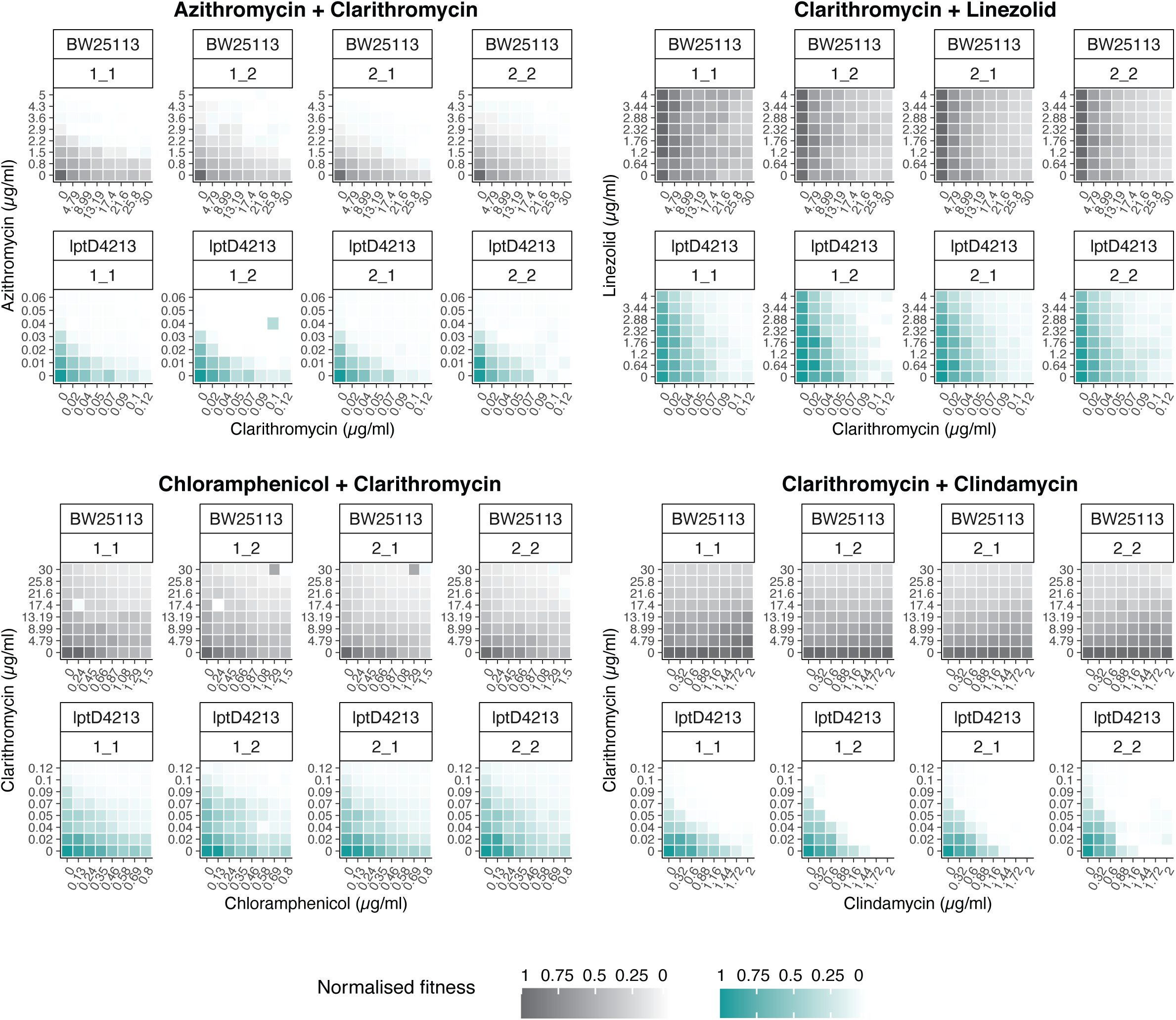
Protein synthesis inhibitors interact neutrally in wild-type *E. coli* BW25113 and synergistically in *E. coli lptD4213* mutant. Checkerboards from which Bliss interaction scores shown in Fig. 3c were derived. Combinations were tested in each strain in two biological and two technical replicates. Synergy, green; Neutrality, grey.

**Extended Data Fig. 10.**
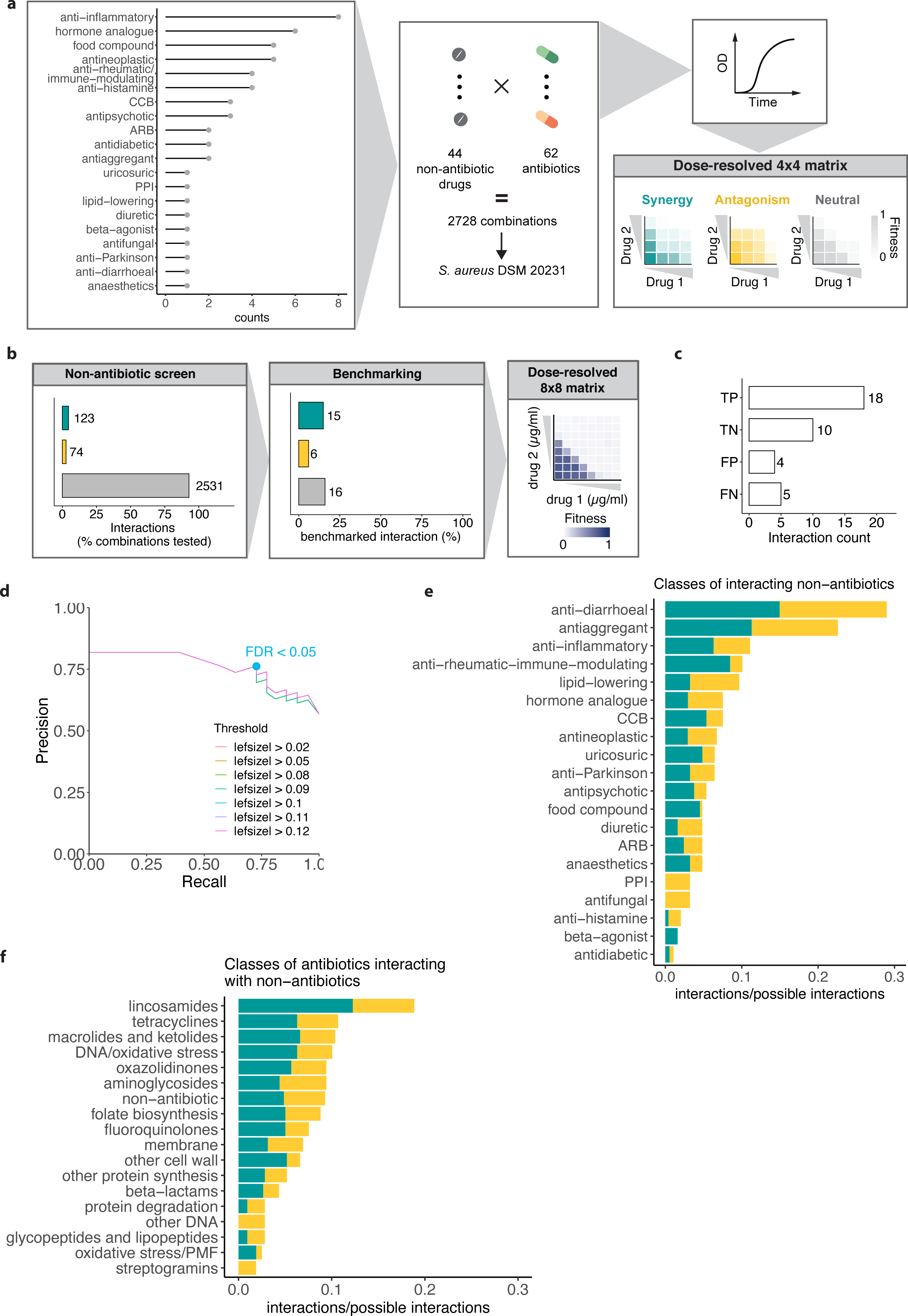
Non-antibiotic drug screen and benchmarking. **a**, Schematic representation of non-antibiotic drug high-throughput screen. 44 drugs belonging to different therapeutic classes (**Supplementary Table 6)** were tested in combination with 62 antibiotics at three concentrations in *S. aureus* DSM 20231. The resulting 2728 combinations were tested in broth microdilution (Methods). **b**, 37 drug combinations (**Supplementary Table 6**) were selected for benchmarking and tested in 8 x 8 concentration checkerboards (Methods). **c**, True-positive (TP), true-negative (TN), false-positive (FP) and false-negative (FN) abundance in the benchmarking set for optimal thresholds shown in **d**. **d**, Screen precision and recall against the benchmarking set are assessed for different effect-size thresholds. Precision-recall curves are shown for FDR intervals ranging from 0 to 1, increasing by 0.005. The chosen significance value for the screen (FDR < 0.05) is highlighted for the effect-size curve (|0.1|) providing best precision and recall. **e-f**, Interaction abundance for classes of non-antibiotics (**e**) and classes of antibiotics (**f**). Interactions detected in *S. aureus* DSM 20231 between antibiotics classes and 52 non-antibiotics tested in both the original screen (n = 8) and the extended screen (n = 44) were considered (n = 245). PMF, Proton-motive force; CCB, calcium-channel blocker; PPI, proton-pump inhibitors; ARB, angiotensin-receptor blocker. Synergies are depicted in green, antagonisms in yellow.

**Extended Data Fig. 11.**
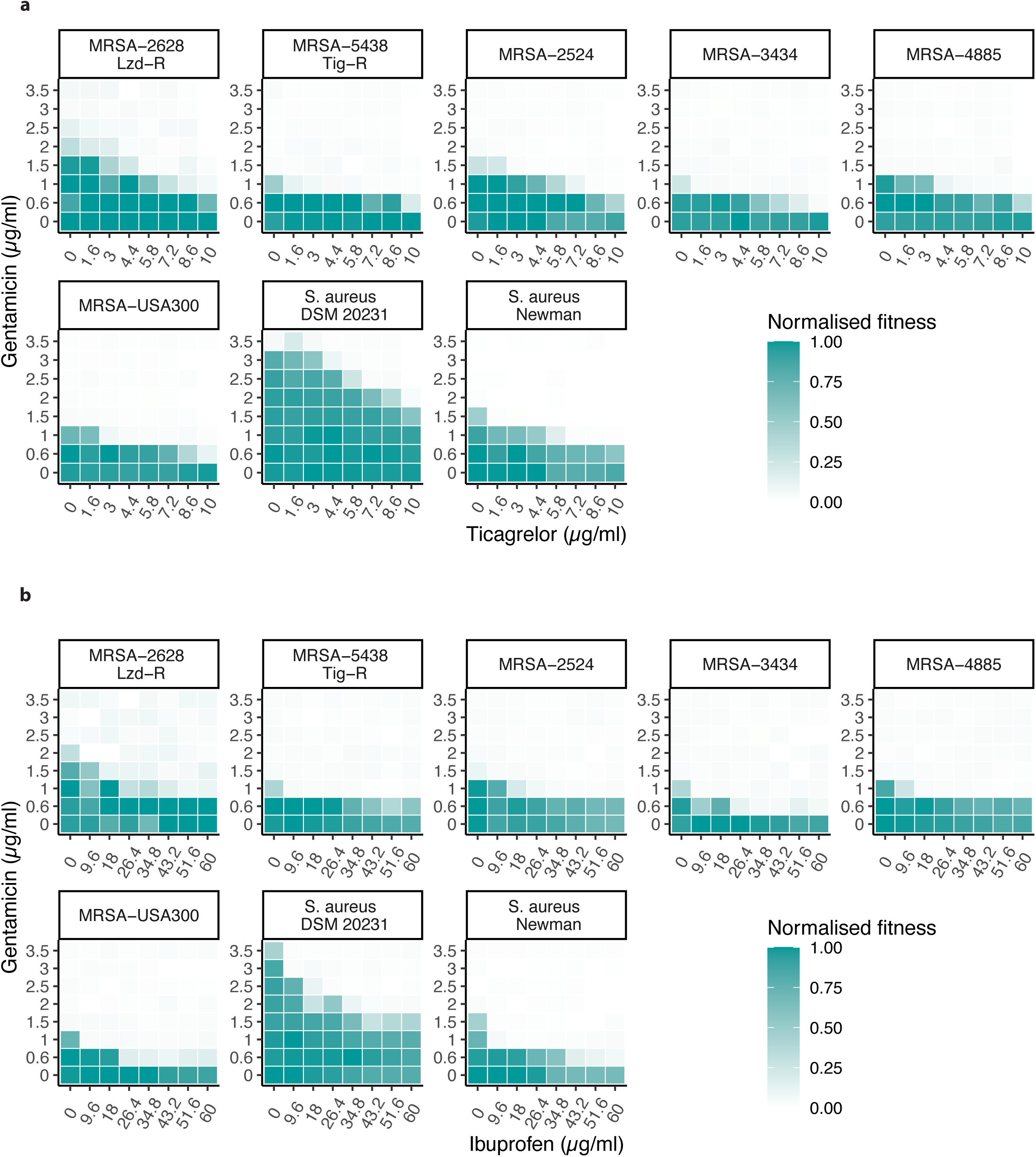
Synergies between antibiotics and non-antibiotic drugs detected in model S. aureus strains are also effective in MRSA clinical isolates. Gentamicin synergies with ibuprofen (a) and ticagrelor (b) were tested in 8 x 8 checkerboards in model MSSA (Newman and DSM 20231) and MRSA (USA300) strains, and in 5 clinical MRSA strains from different clonal complexes, isolated from different infection sites, and bearing different resistance profiles (**Supplementary Table 1**). Results are obtained and represented as in Fig. 2d. One representative checkerboard of n = 2 (biological replicates) is shown here (for the second replicate see Supplementary File). Lzd-R, linezolid-resistant; Tig-R, tigecycline-resistant.

**Extended Data Fig. 12.**
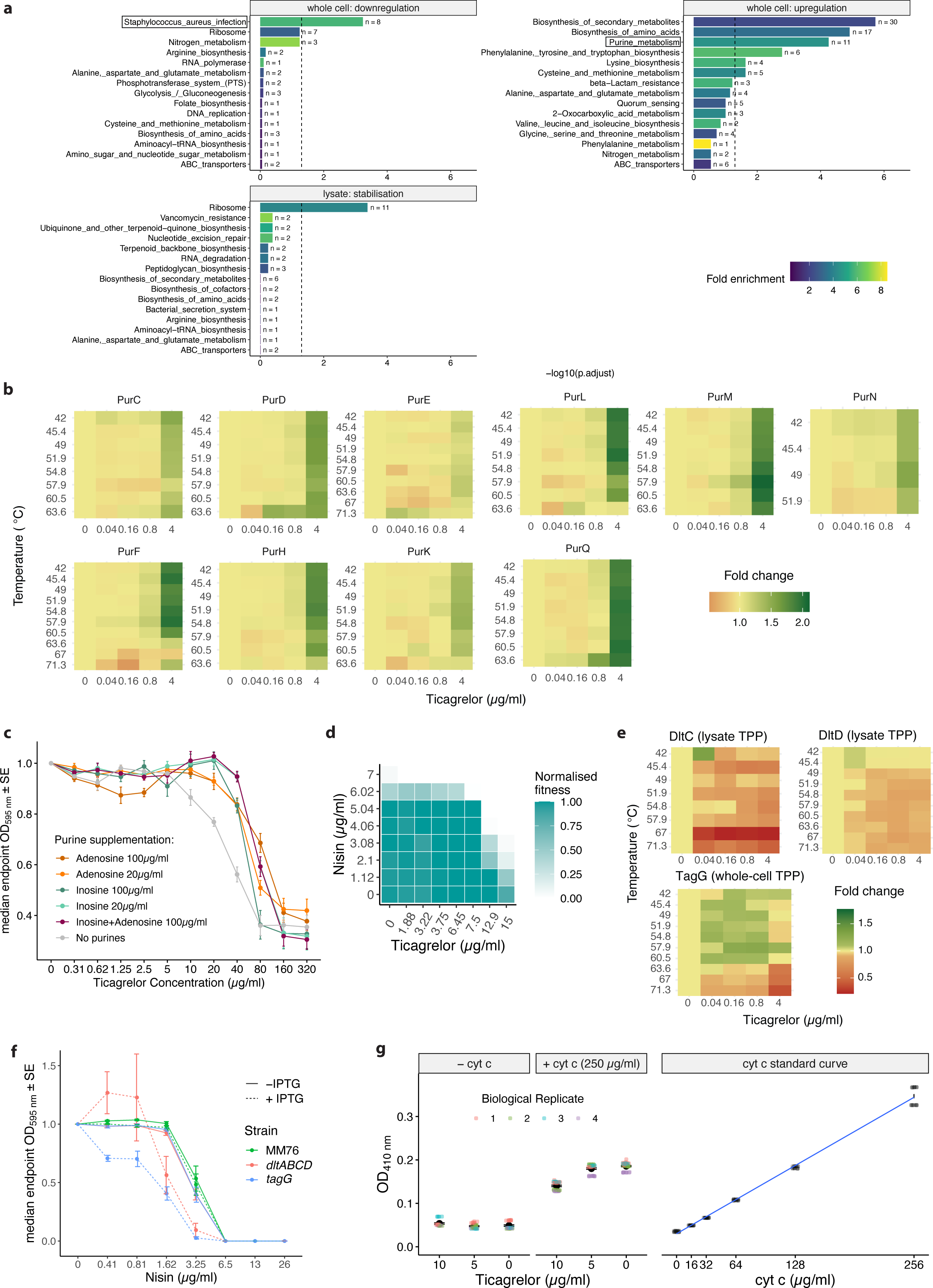
Ticagrelor affects purine and teichoic acid biosynthesis. **a**, KEGG enrichment of hits at 5% FDR from whole-cell or lysate samples. Only sets yielding significant enrichments (whole-cell down- and up-regulation, and lysate stabilisation) are shown (**Supplementary Table 7**). The first 15 terms in order of significance are shown. The dashed lines mark the enrichment significance cut-off (adjusted p-value < 0.05, Fisher’s exact test). The number of protein hits is annotated for each term. **b**, Thermal stability profiles of members of the purine biosynthesis pathway. Protein fold change is shown for each temperature and ticagrelor concentration. **c**, Growth (endpoint-OD_595nm_ after 11h, corresponding to the beginning of stationary phase for the untreated control, Methods) measured in the presence of serial two-fold dilutions of ticagrelor in presence or absence of purines at the indicated concentration, normalised by no-drug controls, in *S. aureus* Newman in SSM9PR defined medium (median across four biological replicates; error bars represent standard error; Methods). **d**, Ticagrelor synergizes with nisin *in vitro*. Results are obtained and represented as in Fig. 2d. **e**, Thermal stability profiles of proteins involved in teichoic acid biosynthesis, represented as in **b**. **f**, Growth (endpoint OD_595nm_, corresponding to the beginning of stationary phase for the control strain MM76, Methods, **Supplementary File**) measured in the presence of serial two-fold dilutions of nisin, normalised by no-drug controls, in the *S. aureus* IPTG-inducible knockdown mutants *dltABCD* and *tagG* and their control strain MM76 (Methods), in presence or absence of 500 µM IPTG to induce maximal knockdown of the gene targeted (median across four biological replicates; error bars represent standard error). All strains are grown in presence of 5 µg/ml erythromycin and 10 µg/ml chloramphenicol to maintain the CRISPRi plasmids^72^ (Methods). For all controls and full growth curves see Supplementary File. **g**, Raw data (OD_410nm_) from Fig. 5g are shown alongside all controls (samples not incubated with 250 µg/ml cytochrome c, cytochrome c standard curve including buffer control). The linear fit for the cytochrome c standard curve used to infer the unbound cytochrome C fraction in supernatants is shown (n = 4, mean and standard error of the mean are shown. Data points represent reads (n=4 biological replicates for each condition, Methods).

## Supplementary File

This file contains the heckerboards in **ED Fig. 7a-b** and **ED Fig. 11a-b** and full growth curves and controls for data in **Fig. 5f** and **Extended Data Fig. 12c** and **f**.

## References

1. Brochado, A. R., et al. Species-specific activity of antibacterial drug combinations. Nature 559, 259–263 (2018).

2. Eliopoulos, GM, Moellering, RC Jr. Antimicrobial combinations. in *Antibiotics in Laboratory Medicine*, Fourth Edition (ed. Lorian, V.) 330–396 (Williams & Wilkins, 1996).

3. Murray, C. J. L., et al. Global burden of bacterial antimicrobial resistance in 2019: a systematic analysis. Lancet 399, 629–655 (2022).

4. Cook, M. A. & Wright, G. D. The past, present, and future of antibiotics. Sci. Transl. Med. 14, eabo7793 (2022).

5. Tyers, M. & Wright, G. D. Drug combinations: a strategy to extend the life of antibiotics in the 21st century. Nat. Rev. Microbiol. 17, 141–155 (2019).

6. Ejim, L., et al. Combinations of antibiotics and nonantibiotic drugs enhance antimicrobial efficacy. Nat. Chem. Biol. 7, 348–350 (2011).

7. Taylor, P. L., Rossi, L., De Pascale, G. & Wright, G. D. A forward chemical screen identifies antibiotic adjuvants in Escherichia coli. ACS Chem. Biol. 7, 1547–1555 (2012).

8. Farha, M. A., Verschoor, C. P., Bowdish, D. & Brown, E. D. Collapsing the proton motive force to identify synergistic combinations against Staphylococcus aureus. Chem. Biol. 20, 1168–1178 (2013).

9. Farha, M. A., et al. Inhibition of WTA synthesis blocks the cooperative action of PBPs and sensitizes MRSA to β-lactams. ACS Chem. Biol. 8, 226–233 (2013).

10. Campbell, J., et al. Synthetic lethal compound combinations reveal a fundamental connection between wall teichoic acid and peptidoglycan biosyntheses in Staphylococcus aureus. ACS Chem. Biol. 6, 106–116 (2011).

11. Lázár, V., Snitser, O., Barkan, D. & Kishony, R. Antibiotic combinations reduce Staphylococcus aureus clearance. Nature 610, 540–546 (2022).

12. Jawetz, E. & Gunnison, J. B. Studies on antibiotic synergism and antagonism: a scheme of combined antibiotic action. Antibiot. Chemother. 2, 243–248 (1952).

13. Roemhild, R., Bollenbach, T. & Andersson, D. I. The physiology and genetics of bacterial responses to antibiotic combinations. Nat. Rev. Microbiol. 20, 478–490 (2022).

14. Kantor, E. D., Rehm, C. D., Haas, J. S., Chan, A. T. & Giovannucci, E. L. Trends in Prescription Drug Use Among Adults in the United States From 1999-2012. JAMA 314, 1818–1831 (2015).

15. Centers for Disease Control and Prevention (CDC). National Center for Health Statistics (NCHS). National Health and Nutrition Examination Survey Data. Hyattsville, MD: U.S. Department of Health and Human Services, Centers for Disease Control and Prevention, 2021, https://wwwn.cdc.gov/nchs/nhanes/2017-2018/p_rxq_rx.htm.

16. Pai, M. P., Momary, K. M. & Rodvold, K. A. Antibiotic drug interactions. Med. Clin. North Am. 90, 1223–1255 (2006).

17. Tacconelli, E., et al. Discovery, research, and development of new antibiotics: the WHO priority list of antibiotic-resistant bacteria and tuberculosis. Lancet Infect. Dis. 18, 318– 327 (2018).

18. Farha, M. A., et al. Antagonism screen for inhibitors of bacterial cell wall biogenesis uncovers an inhibitor of undecaprenyl diphosphate synthase. Proc. Natl. Acad. Sci. U. S. A. 112, 11048–11053 (2015).

19. Bliss, C. I. THE TOXICITY OF POISONS APPLIED JOINTLY1. Ann. Appl. Biol. 26, 585– 615 (1939).

20. Jawetz, E. The use of combinations of antimicrobial drugs. Annu. Rev. Pharmacol. 8, 151–170 (1968).

21. Mende, D. R., Sunagawa, S., Zeller, G. & Bork, P. Accurate and universal delineation of prokaryotic species. Nat. Methods 10, 881–884 (2013).

22. Dillon, N., et al. Surprising synergy of dual translation inhibition vs. Acinetobacter baumannii and other multidrug-resistant bacterial pathogens. EBioMedicine 46, 193–201 (2019).

23. Bollenbach, T., Quan, S., Chait, R. & Kishony, R. Nonoptimal microbial response to antibiotics underlies suppressive drug interactions. Cell 139, 707–718 (2009).

24. Tang, H.-J., et al. Cephalosporin-Glycopeptide Combinations for Use against Clinical Methicillin-Resistant Staphylococcus aureus Isolates: Enhanced In vitro Antibacterial Activity. Front. Microbiol. 8, 884 (2017).

25. Lai, C.-C., Chen, C.-C., Chuang, Y.-C. & Tang, H.-J. Combination of cephalosporins with vancomycin or teicoplanin enhances antibacterial effect of glycopeptides against heterogeneous vancomycin-intermediate Staphylococcus aureus (hVISA) and VISA. Sci. Rep. 7, 41758 (2017).

26. Rieg, S., et al. Combination antimicrobial therapy in patients with Staphylococcus aureus bacteraemia-a post hoc analysis in 964 prospectively evaluated patients. Clin. Microbiol. Infect. 23, 406.e1–406.e8 (2017).

27. Leone, S., Noviello, S. & Esposito, S. Combination antibiotic therapy for the treatment of infective endocarditis due to enterococci. Infection 44, 273–281 (2016).

28. Baddour, L. M., et al. Combination antibiotic therapy lowers mortality among severely ill patients with pneumococcal bacteremia. Am. J. Respir. Crit. Care Med. 170, 440–444 (2004).

29. Habib, G., et al. 2015 ESC Guidelines for the management of infective endocarditis: The Task Force for the Management of Infective Endocarditis of the European Society of Cardiology (ESC). Endorsed by: European Association for Cardio-Thoracic Surgery (EACTS), the European Association of Nuclear Medicine (EANM). Eur. Heart J. 36, 3075– 3128 (2015).

30. Bartash, R. & Nori, P. Beta-lactam combination therapy for the treatment of Staphylococcus aureus and Enterococcus species bacteremia: A summary and appraisal of the evidence. Int. J. Infect. Dis. 63, 7–12 (2017).

31. Ida, T., et al. Antagonism between aminoglycosides and beta-lactams in a methicillin-resistant Staphylococcus aureus isolate involves induction of an aminoglycoside-modifying enzyme. Antimicrob. Agents Chemother. 46, 1516–1521 (2002).

32. Vakulenko, S. B. & Mobashery, S. Versatility of aminoglycosides and prospects for their future. Clin. Microbiol. Rev. 16, 430–450 (2003).

33. Paul, M., Lador, A., Grozinsky-Glasberg, S. & Leibovici, L. Beta lactam antibiotic monotherapy versus beta lactam-aminoglycoside antibiotic combination therapy for sepsis. Cochrane Database Syst. Rev. CD003344 (2014).

34. Typas, A. & Sourjik, V. Bacterial protein networks: properties and functions. Nat. Rev. Microbiol. 13, 559–572 (2015).

35. Kavčič, B., Tkačik, G. & Bollenbach, T. Mechanisms of drug interactions between translation-inhibiting antibiotics. Nat. Commun. 11, 4013 (2020).

36. Sampson, B. A., Misra, R. & Benson, S. A. Identification and characterization of a new gene of Escherichia coli K-12 involved in outer membrane permeability. Genetics 122, 491–501 (1989).

37. Ruiz, N., Falcone, B., Kahne, D. & Silhavy, T. J. Chemical conditionality: a genetic strategy to probe organelle assembly. Cell vol. 121 307–317 (2005).

38. Sauvage, E., Kerff, F., Terrak, M., Ayala, J. A. & Charlier, P. The penicillin-binding proteins: structure and role in peptidoglycan biosynthesis. FEMS Microbiol. Rev. 32, 234– 258 (2008).

39. Egan, A. J. F., Errington, J. & Vollmer, W. Regulation of peptidoglycan synthesis and remodelling. Nat. Rev. Microbiol. 18, 446–460 (2020).

40. Schulz, M., Iwersen-Bergmann, S., Andresen, H. & Schmoldt, A. Therapeutic and toxic blood concentrations of nearly 1,000 drugs and other xenobiotics. Crit. Care 16, R136 (2012).

41. Kavanaugh, M. L. & Jerman, J. Contraceptive method use in the United States: trends and characteristics between 2008, 2012 and 2014. Contraception 97, 14–21 (2018).

42. Chan, E. W. L., Yee, Z. Y., Raja, I. & Yap, J. K. Y. Synergistic effect of non-steroidal anti-inflammatory drugs (NSAIDs) on antibacterial activity of cefuroxime and chloramphenicol against methicillin-resistant Staphylococcus aureus. J Glob Antimicrob Resist 10, 70–74 (2017).

43. Zimmermann, P. & Curtis, N. The effect of aspirin on antibiotic susceptibility. Expert Opinion on Therapeutic Targets vol. 22 967–972 Preprint at https://doi.org/10.1080/14728222.2018.1527314 (2018).

44. Cohen, S. P., Levy, S. B., Foulds, J. & Rosner, J. L. Salicylate induction of antibiotic resistance in Escherichia coli: activation of the mar operon and a mar-independent pathway. J. Bacteriol. 175, 7856–7862 (1993).

45. Price, C. T., Lee, I. R. & Gustafson, J. E. The effects of salicylate on bacteria. Int. J. Biochem. Cell Biol. 32, 1029–1043 (2000).

46. Husted, S. & van Giezen, J. J. J. Ticagrelor: the first reversibly binding oral P2Y12 receptor antagonist. Cardiovasc. Ther. 27, 259–274 (2009).

47. Storey, R. F., et al. Lower mortality following pulmonary adverse events and sepsis with ticagrelor compared to clopidogrel in the PLATO study. Platelets 25, 517–525 (2014).

48. Sexton, T. R., et al. Ticagrelor Reduces Thromboinflammatory Markers in Patients With Pneumonia. JACC Basic Transl Sci 3, 435–449 (2018).

49. Sun, J., et al. Repurposed drugs block toxin-driven platelet clearance by the hepatic Ashwell-Morell receptor to clear *Staphylococcus aureus* bacteremia. Sci. Transl. Med. 13, (2021).

50. Ulloa, E. R., Uchiyama, S., Gillespie, R., Nizet, V. & Sakoulas, G. Ticagrelor Increases Platelet-Mediated Staphylococcus aureus Killing, Resulting in Clearance of Bacteremia. J. Infect. Dis. 224, 1566–1569 (2021).

51. Lancellotti, P., et al. Antibacterial Activity of Ticagrelor in Conventional Antiplatelet Dosages Against Antibiotic-Resistant Gram-Positive Bacteria. JAMA Cardiol 4, 596–599 (2019).

52. Becher, I., et al. Thermal profiling reveals phenylalanine hydroxylase as an off-target of panobinostat. Nat. Chem. Biol. 12, 908–910 (2016).

53. Mateus, A., et al. Thermal proteome profiling in bacteria: probing protein state in vivo. Mol. Syst. Biol. 14, e8242 (2018).

54. Mateus, A., et al. Thermal proteome profiling for interrogating protein interactions. Mol. Syst. Biol. 16, e9232 (2020).

55. Cheng, A. G., et al. Contribution of coagulases towards Staphylococcus aureus disease and protective immunity. PLoS Pathog. 6, e1001036 (2010).

56. Taber, H. W., Mueller, J. P., Miller, P. F. & Arrow, A. S. Bacterial uptake of aminoglycoside antibiotics. Microbiol. Rev. 51, 439–457 (1987).

57. Wood, B. M., Santa Maria, J. P., Jr, Matano, L. M., Vickery, C. R. & Walker, S. A partial reconstitution implicates DltD in catalyzing lipoteichoic acid d-alanylation. J. Biol. Chem. 293, 17985–17996 (2018).

58. Xia, G., Kohler, T. & Peschel, A. The wall teichoic acid and lipoteichoic acid polymers of Staphylococcus aureus. Int. J. Med. Microbiol. 300, 148–154 (2010).

59. Pasquina, L., et al. A synthetic lethal approach for compound and target identification in Staphylococcus aureus. Nat. Chem. Biol. 12, 40–45 (2016).

60. Brown, S., Santa Maria, J. P., Jr & Walker, S. Wall teichoic acids of gram-positive bacteria. Annu. Rev. Microbiol. 67, 313–336 (2013).

61. Sastry, S. & Doi, Y. Fosfomycin: Resurgence of an old companion. J. Infect. Chemother. 22, 273–280 (2016).

62. Ericsson, C. D., DuPont, H. L., Okhuysen, P. C., Jiang, Z.-D. & DuPont, M. W. Loperamide plus azithromycin more effectively treats travelers’ diarrhea in Mexico than azithromycin alone. J. Travel Med. 14, 312–319 (2007).

63. Miró-Canturri, A., Ayerbe-Algaba, R. & Smani, Y. Drug Repurposing for the Treatment of Bacterial and Fungal Infections. Front. Microbiol. 10, 41 (2019).

64. Phanchana, M., et al. Repurposing a platelet aggregation inhibitor ticagrelor as an antimicrobial against Clostridioides difficile. Sci. Rep. 10, 6497 (2020).

65. Maier, L., et al. Unravelling the collateral damage of antibiotics on gut bacteria. Nature (2021) doi:10.1038/s41586-021-03986-2.

66. Kunst, F., et al. The complete genome sequence of the gram-positive bacterium Bacillus subtilis. Nature 390, 249–256 (1997).

67. Slager, J., Aprianto, R. & Veening, J.-W. Deep genome annotation of the opportunistic human pathogen Streptococcus pneumoniae D39. Nucleic Acids Res. 46, 9971–9989 (2018).

68. Baba, T., Bae, T., Schneewind, O., Takeuchi, F. & Hiramatsu, K. Genome Sequence of Staphylococcus aureus Strain Newman and Comparative Analysis of Staphylococcal Genomes: Polymorphism and Evolution of Two Major Pathogenicity Islands. J. Bacteriol. 190, 300–310 (2008).

69. Shiroma, A., et al. First Complete Genome Sequences of Staphylococcus aureus subsp. aureus Rosenbach 1884 (DSM 20231T), Determined by PacBio Single-Molecule Real-Time Technology. Genome Announc. 3, (2015).

70. Martin B, García P, Castanié M-P, Claverys J-P. The recA gene of Streptococcus pneumoniae is part of acompetence-induced operon and controls lysogenic induction. Molecular Microbiology 15, 367–379 (1995).

71. Reed, P., et al. Staphylococcus aureus Survives with a Minimal Peptidoglycan Synthesis Machine but Sacrifices Virulence and Antibiotic Resistance. PLoS Pathog. 11, e1004891 (2015).

72. Stamsås, G. A., et al. CozEa and CozEb play overlapping and essential roles in controlling cell division in Staphylococcus aureus. Mol. Microbiol. 109, 615–632 (2018).

73. Huber, P. J. Robust Estimation of a Location Parameter. Ann. Math. Stat. 35, 73–101 (1964).

74. Goldoni, M. & Johansson, C. A mathematical approach to study combined effects of toxicants in vitro: evaluation of the Bliss independence criterion and the Loewe additivity model. Toxicol. In Vitro 21, 759–769 (2007).

75. R Core Team. R: A language and environment for statistical computing. R Foundation for Statistical Computing, Vienna, Austria. URL https://www.r-project.org/. (2021).

76. RStudio Team. RStudio: Integrated Development Environment for R. RStudio, PBC, Boston, MA URL http://www.rstudio.com/. (2021).

77. Shannon, P., et al. Cytoscape: a software environment for integrated models of biomolecular interaction networks. Genome Res. 13, 2498–2504 (2003).

78. Bajusz, D., Rácz, A. & Héberger, K. Why is Tanimoto index an appropriate choice for fingerprint-based similarity calculations? J. Cheminform. 7, 20 (2015).

79. Morgan, H. L. The Generation of a Unique Machine Description for Chemical Structures-A Technique Developed at Chemical Abstracts Service. J. Chem. Doc. 5, 107–113 (1965).

80. Milanese, A., et al. Microbial abundance, activity and population genomic profiling with mOTUs2. Nat. Commun. 10, 1014 (2019).

81. Rognes, T., Flouri, T., Nichols, B., Quince, C. & Mahé, F. VSEARCH: a versatile open source tool for metagenomics. PeerJ 4, e2584 (2016).

82. Edgar, R. C. MUSCLE: multiple sequence alignment with high accuracy and high throughput. Nucleic Acids Res. 32, 1792–1797 (2004).

83. Guindon, S., et al. New algorithms and methods to estimate maximum-likelihood phylogenies: assessing the performance of PhyML 3.0. Syst. Biol. 59, 307–321 (2010).

84. Kurzawa N, Franken H, Anders S, Huber W, Savitski M. TPP2D: Detection of ligand-protein interactions from 2D thermal profiles (DLPTP). R package version 1.4.1, http://bioconductor.org/packages/tpp2d. (2020).

85. Kurzawa, N., et al. Computational analysis of ligand dose range thermal proteome profiles. 2020.05.08.083709 (2020) doi:10.1101/2020.05.08.083709.

86. Kurzawa, N., et al. A computational method for detection of ligand-binding proteins from dose range thermal proteome profiles. Nat. Commun. 11, 5783 (2020).

87. Kanehisa, M. & Goto, S. KEGG: kyoto encyclopedia of genes and genomes. Nucleic Acids Res. 28, 27–30 (2000).

88. Tenenbaum D, M. B. KEGGREST: Client-side REST access to the Kyoto Encyclopedia of Genes and Genomes (KEGG). R package version 1.34.0. (2021).

89. Geistlinger, L., Csaba, G. & Zimmer, R. Bioconductor’s EnrichmentBrowser: seamless navigation through combined results of set- & network-based enrichment analysis. BMC Bioinformatics 17, 45 (2016).

90. Wu, T., et al. clusterProfiler 4.0: A universal enrichment tool for interpreting omics data. Innovation (N Y) 2, 100141 (2021).

91. Radlinski, L. C., et al. Chemical Induction of Aminoglycoside Uptake Overcomes Antibiotic Tolerance and Resistance in Staphylococcus aureus. Cell Chem Biol 26, 1355–1364.e4 (2019).

92. Brötz-Oesterhelt, H. & Vorbach, A. Reprogramming of the Caseinolytic Protease by ADEP Antibiotics: Molecular Mechanism, Cellular Consequences, Therapeutic Potential. Front Mol Biosci 8, 690902 (2021).

93. Weinandy, F., et al. A β-lactone-based antivirulence drug ameliorates Staphylococcus aureus skin infections in mice. ChemMedChem 9, 710–713 (2014).

