## Supplementary File for "High-throughput profiling of drug interactions in Gram-positive bacteria"

**Source Fig. 2d, ED Fig. 7a-b**

**a**

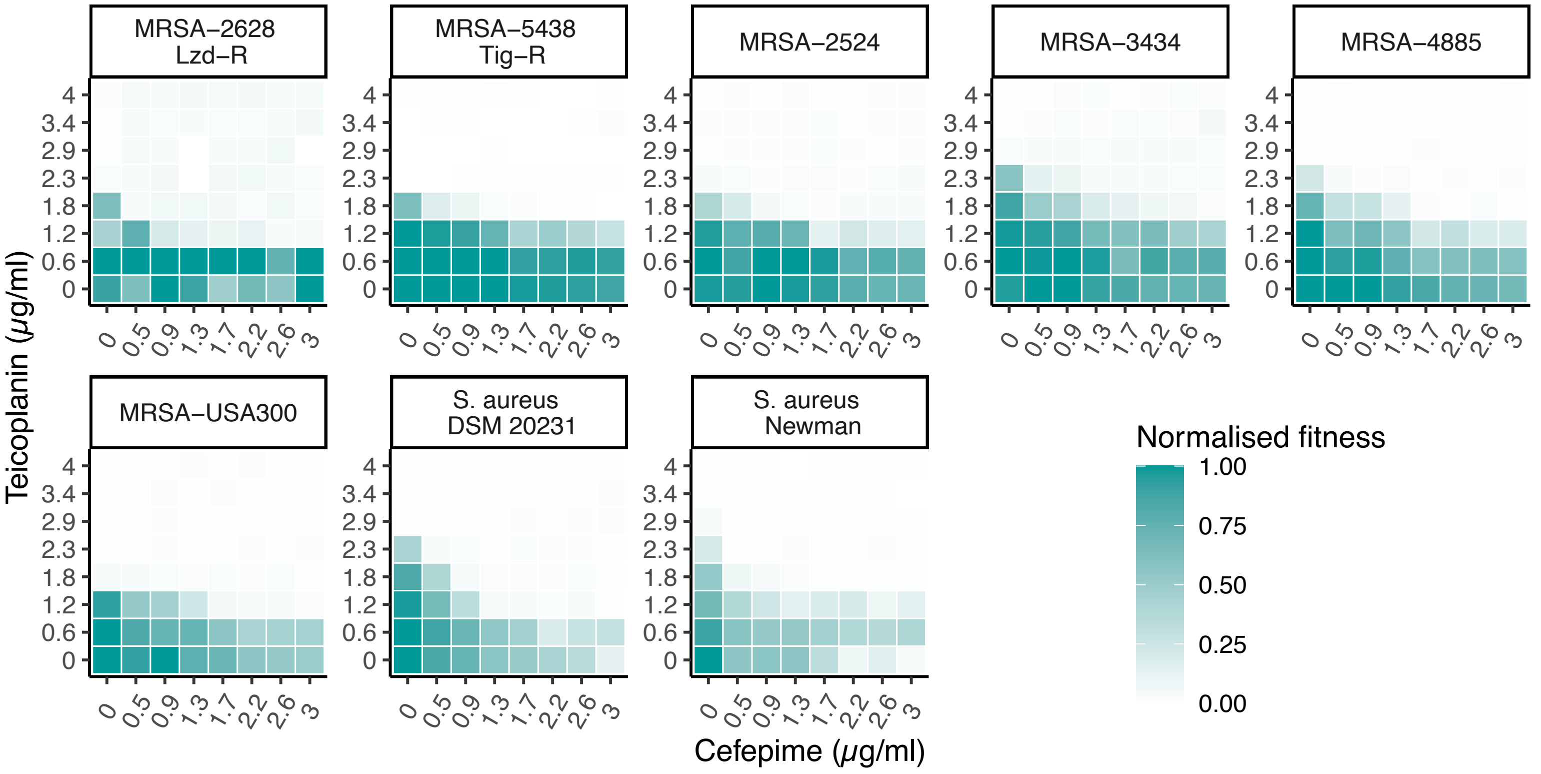


**b**

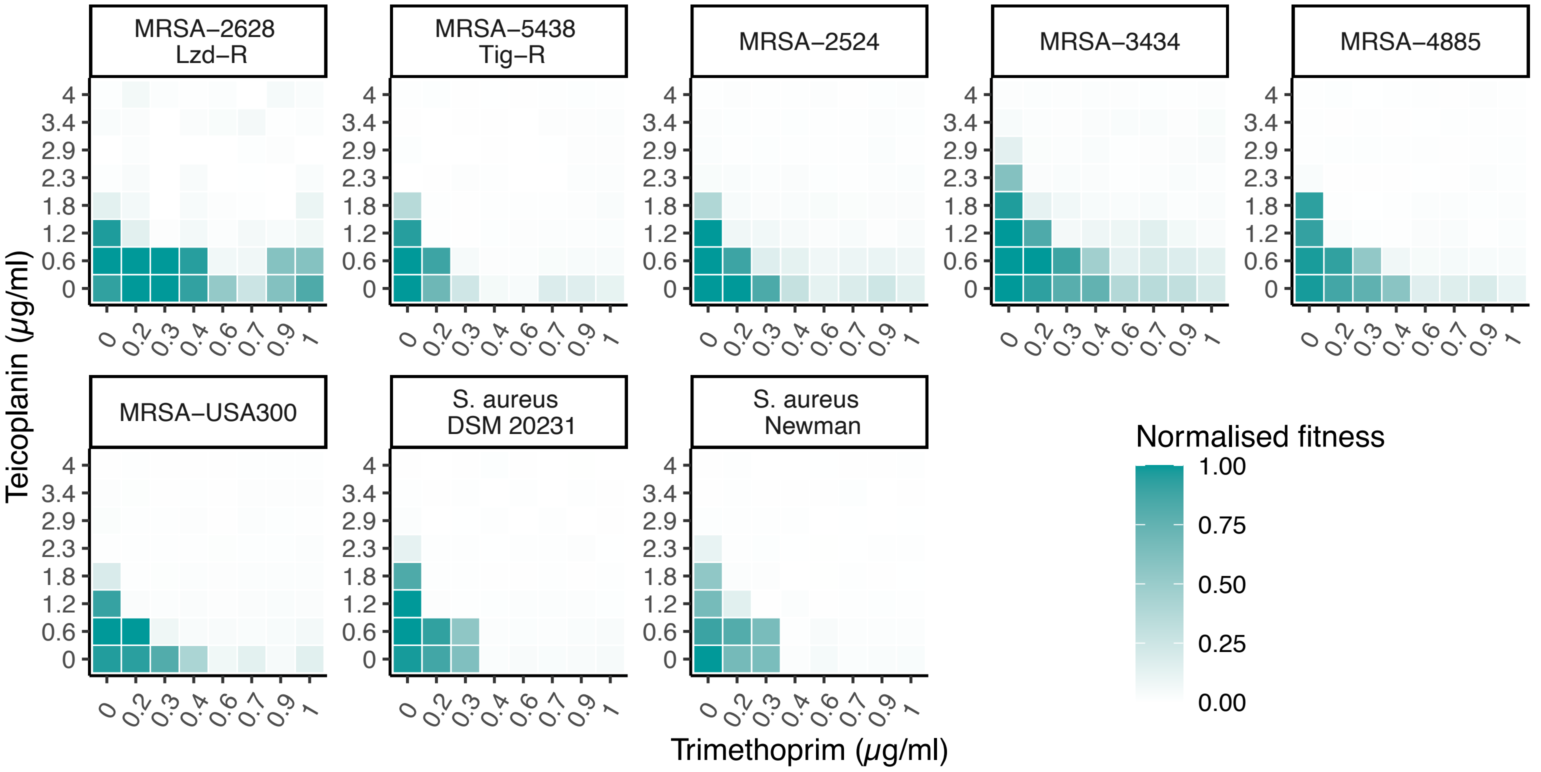


Second biological replicate of two (first shown in Extended Data Fig. 7a-b) for checkerboard assays displayed in Fig. 2d. Results are obtained and represented as in Fig. 2d.

**Source Fig. 4e, Source Fig. 5c, ED Fig. 11a-b**

**a**

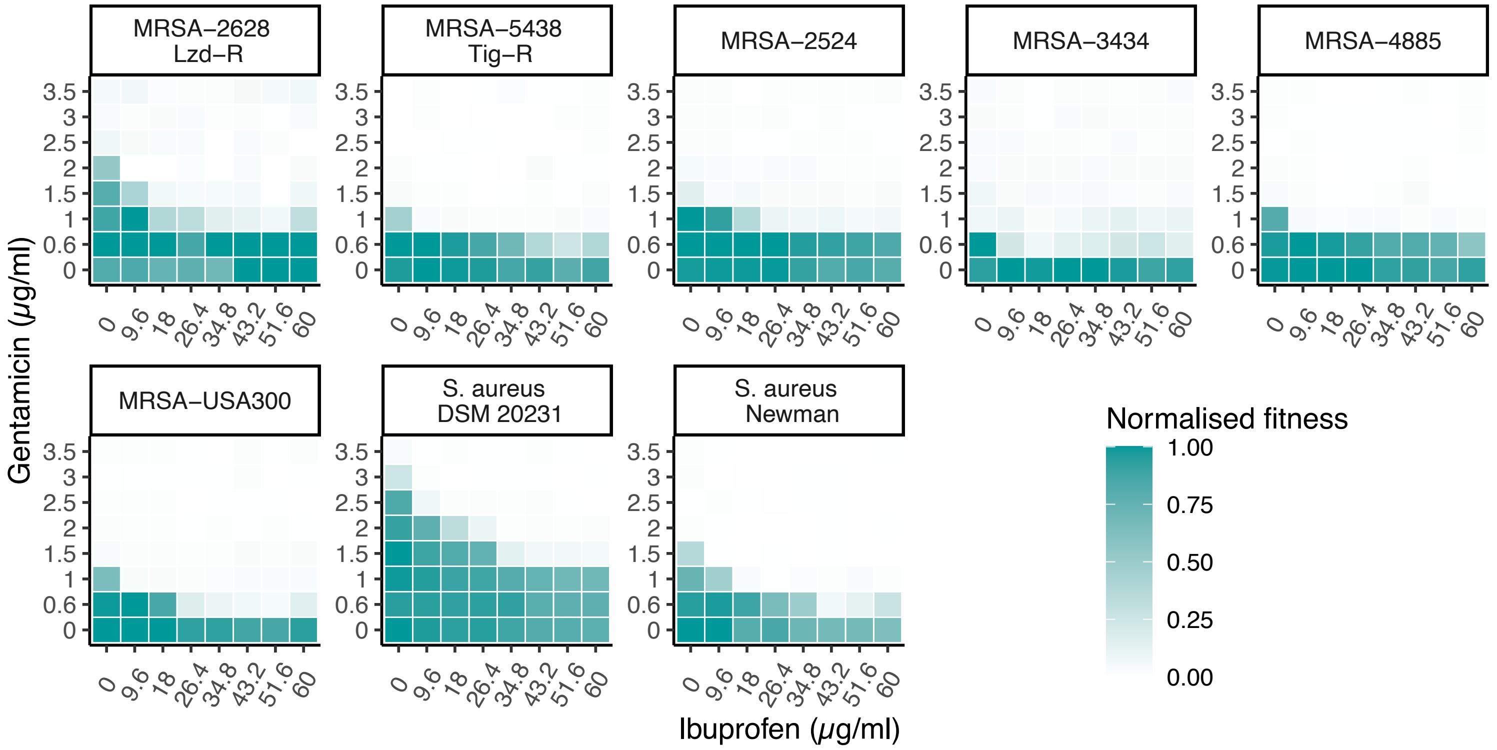


**b**

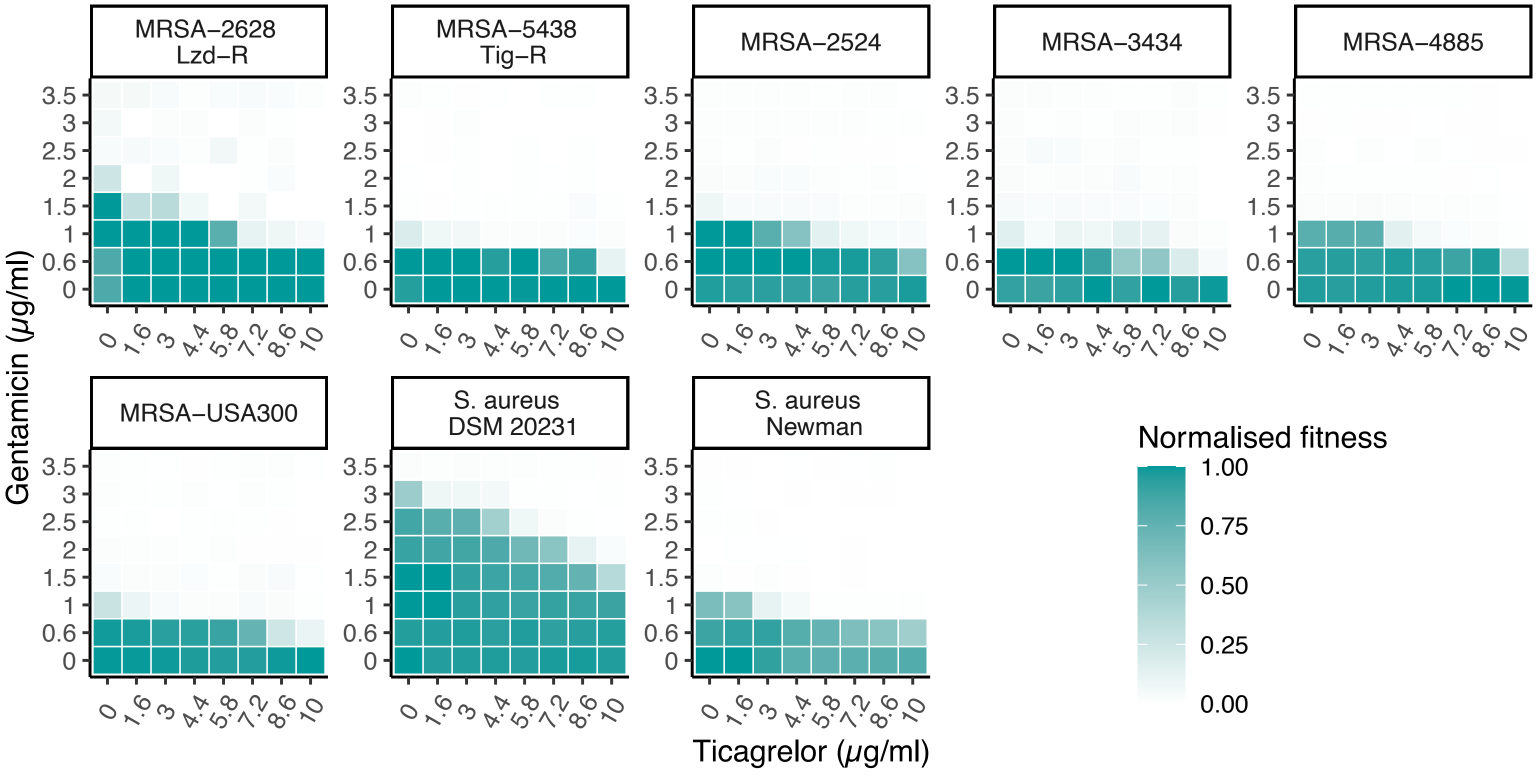


Second biological replicate of two (first shown in Extended Data Fig. 11a-b) for checkerboard assays displayed in Fig. 4e and Fig. 5c, respectively. Results are obtained and represented as in Fig. 2d.

### Source Fig. 5f, ED Fig. 12f

**a**

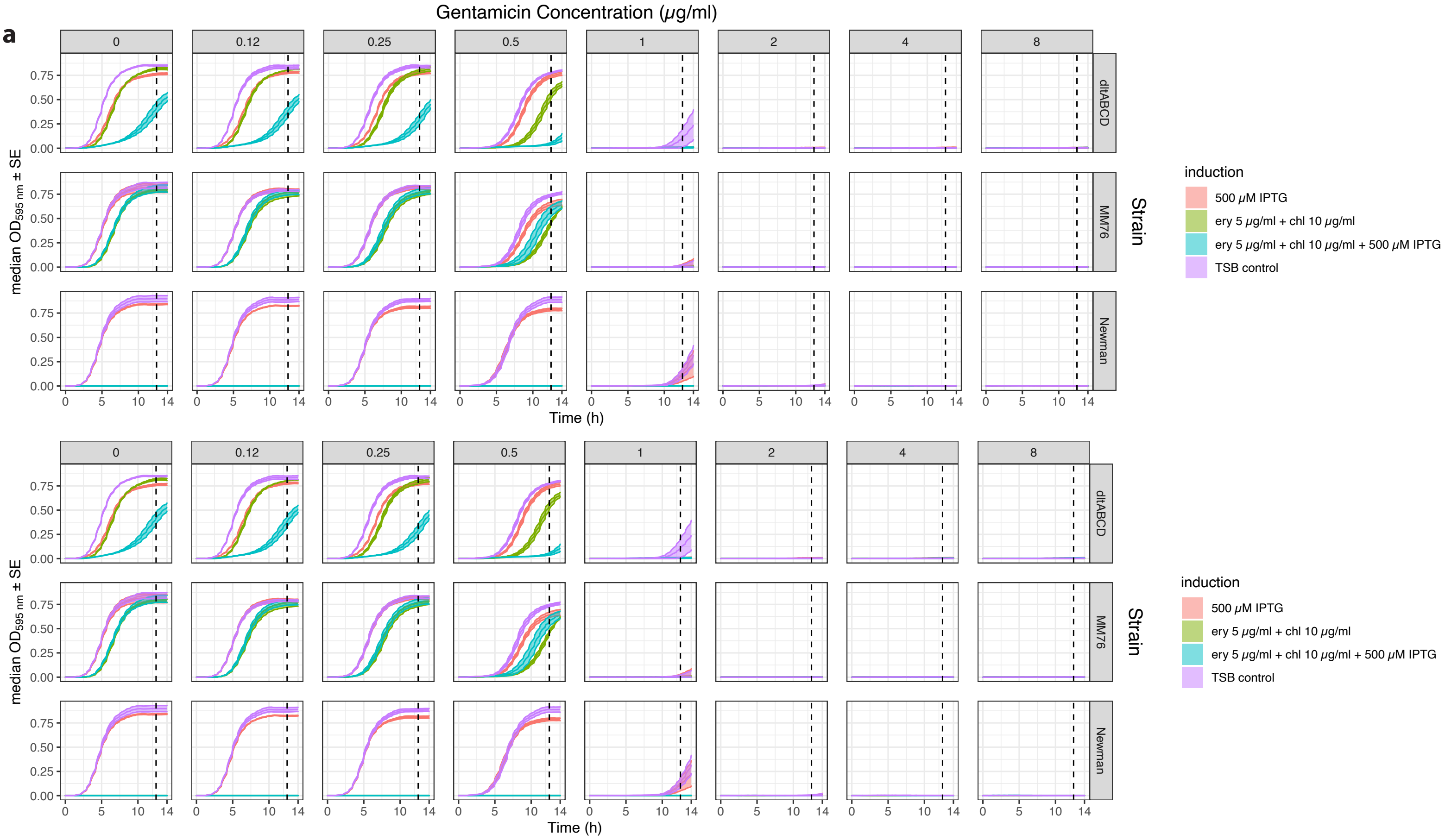


**b**

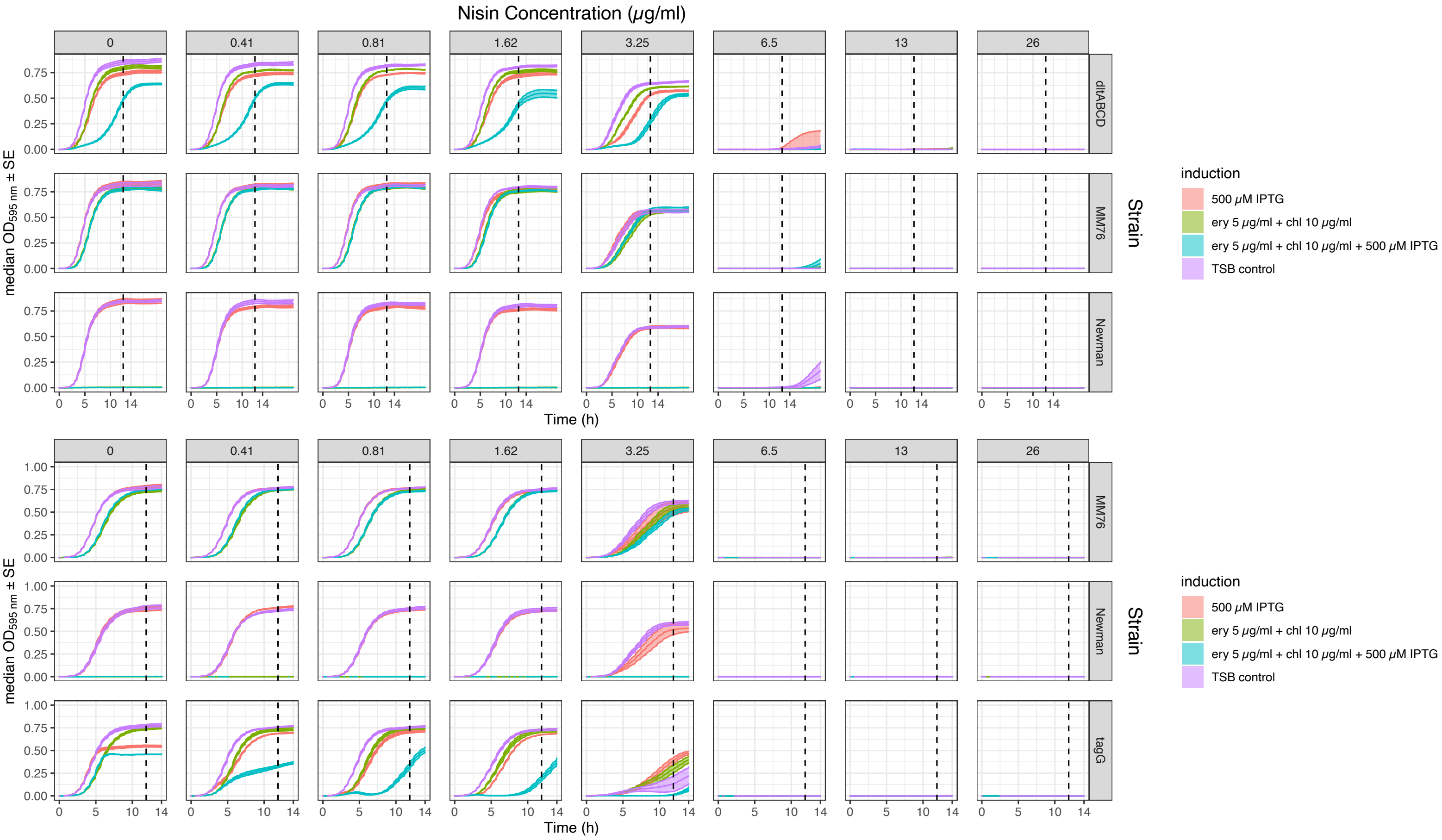


Full growth curves from which dose-response curves depicted in Fig. 5f (**a**) and Extended Data Fig. 12f (**b**) were obtained. Median $OD_{595nm}$ and standard error across four biological replicates are shown. Dashed lines represent the automatically-detected time point of $OD_{595nm}$ plateau for the control strain MM76 upon full induction, which was used to obtain the dose-response curves for each experiment. Ery, erythromycin; chl, chloramphenicol; TSB, Tryptic Soy Broth.

### Source Fig. 5f, ED Fig. 12f

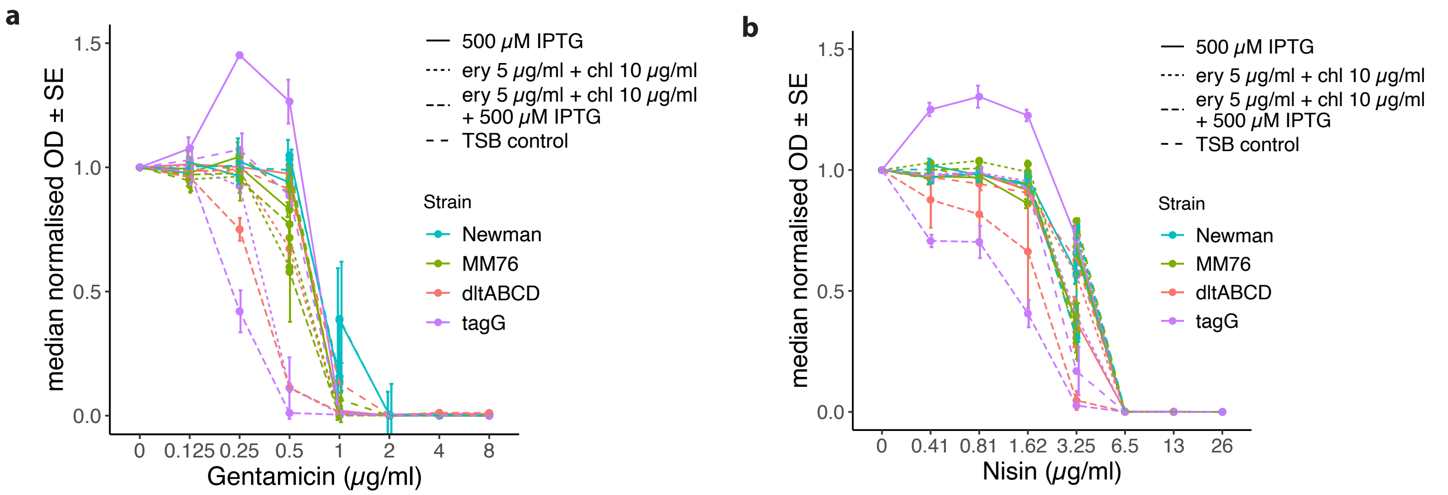


Dose-response curves depicted in Fig. 5f (**a**) and Extended Data Fig. 12f (**b**) complete of all controls. Median $OD_{595nm}$ and standard error across four biological replicates are shown. Ery, erythromycin; chl, chloramphenicol; TSB, Tryptic Soy Broth.

### Source ED Fig. 12c

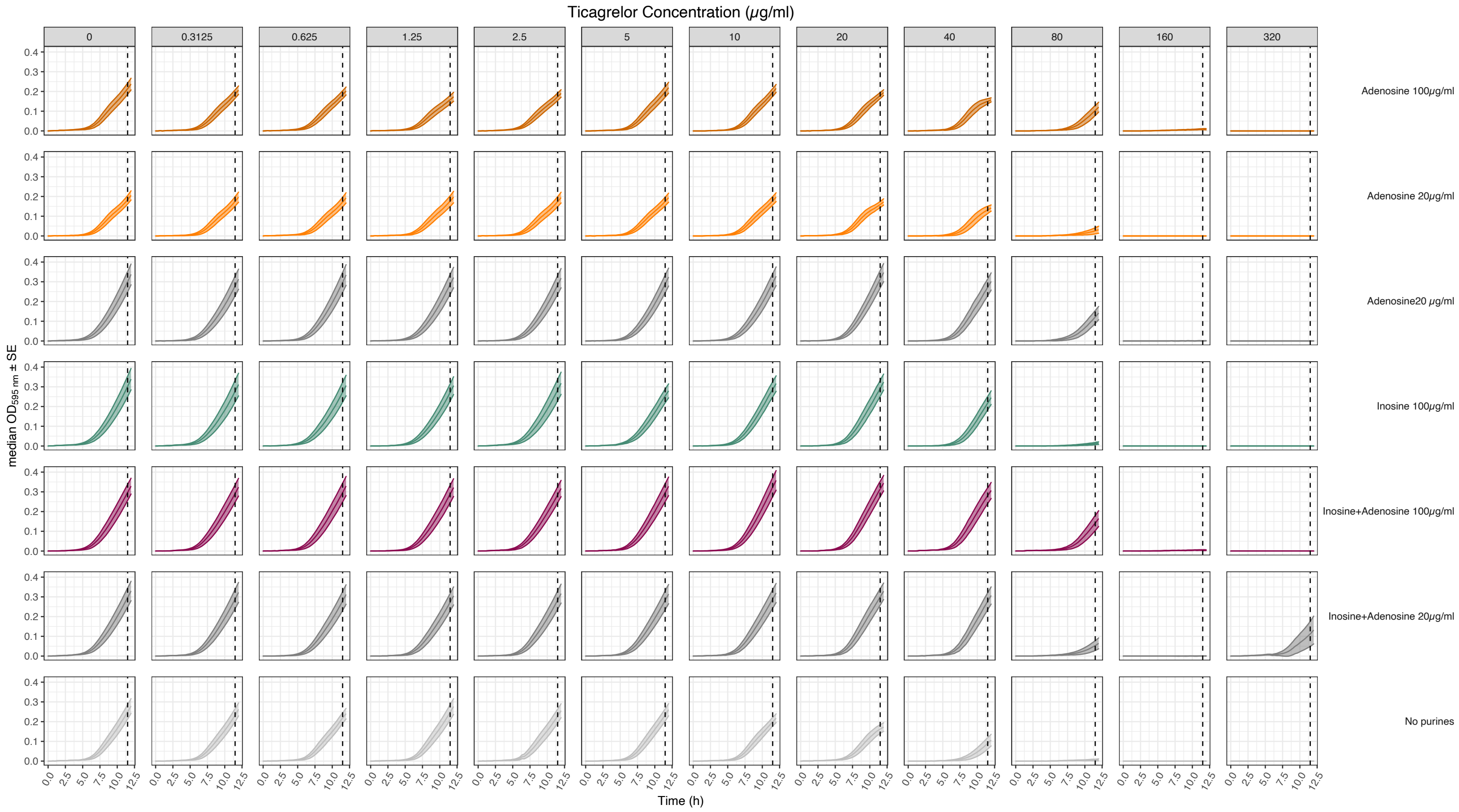


Full growth curves from which dose-response curves depicted in Extended Data Fig. 12c were obtained. Median $OD_{595nm}$ and standard error across four biological replicates are shown. Dashed lines represent the time point used to obtain the dose-response curves. Ery, erythromycin; chl, chloramphenicol; TSB, Tryptic Soy Broth.
